# High-resolution single-particle imaging at 100-200 keV with the Gatan Alpine direct electron detector

**DOI:** 10.1101/2024.02.14.580363

**Authors:** Lieza M. Chan, Brandon J. Courteau, Allison Maker, Mengyu Wu, Benjamin Basanta, Hevatib Mehmood, David Bulkley, David Joyce, Brian C. Lee, Stephen Mick, Sahil Gulati, Gabriel C. Lander, Kliment A. Verba

## Abstract

Developments in direct electron detector technology have played a pivotal role in enabling high-resolution structural studies by cryo-EM at 200 and 300 keV. Yet, theory and recent experiments indicate advantages to imaging at 100 keV, energies for which the current detectors have not been optimized. In this study, we evaluated the Gatan Alpine detector, designed for operation at 100 and 200 keV. Compared to the Gatan K3, Alpine demonstrated a significant DQE improvement at these voltages, specifically a ~4-fold improvement at Nyquist at 100 keV. In single-particle cryo-EM experiments, Alpine datasets yielded better than 2 Å resolution reconstructions of apoferritin at 120 and 200 keV on a ThermoFisher Scientific (TFS) Glacios microscope. We also achieved a ~3.2 Å resolution reconstruction for a 115 kDa asymmetric protein complex, proving its effectiveness with complex biological samples. In-depth analysis revealed that Alpine reconstructions are comparable to K3 reconstructions at 200 keV, and remarkably, reconstruction from Alpine at 120 keV on a TFS Glacios surpassed all but the 300 keV data from a TFS Titan Krios with GIF/K3. Additionally, we show Alpine’s capability for high-resolution data acquisition and screening on lower-end systems by obtaining ~3 Å resolution reconstructions of apoferritin and aldolase at 100 keV and detailed 2D averages of a 55 kDa sample using a side-entry cryo holder. Overall, we show that Gatan Alpine performs well with the standard 200 keV imaging systems and may potentially capture the benefits of lower accelerating voltages, possibly bringing smaller sized particles within the scope of cryo-EM.

## 1. INTRODUCTION

Cryo-electron microscopy (cryo-EM) has become a powerful tool for macromolecular structure determination, providing valuable insights into the mechanisms and functions of proteins and their complexes. In the past two years, the number of structures deposited in the PDB determined by electron microscopy-based methods has reached record-breaking highs, matching nearly 50% of the number of structures determined by x-ray-based methods in the year 2023 (Berman et al. 2000; Protein Data Bank). The number of PDB-deposited cryo-EM structures has rapidly increased each year since the ‘resolution revolution’, driven in great part by technological advancements in both hardware and software, particularly in detector technologies (Kühlbrandt 2014). In general, improvements in electron detectors have led to direct detection (Deptuch et al. 2007; Guerrini et al. 2011), single electron counting (Li, Zheng, et al. 2013; Li, Mooney, et al. 2013), super-resolution modes (Booth 2012; Li, Zheng, et al. 2013), and correlated-double sampling modes (Sun et al. 2021), all of which have dramatically improved the capabilities of cryo-EM. Despite this, cryo-EM still faces challenges, particularly in obtaining structures of biological targets smaller than 50 kDa (Lander and Glaeser 2021) and in being prohibitively expensive (Hand 2020), limiting access to this powerful technique.

Cryo-EM imaging of biological specimens relies on the interaction of electrons with the sample, comprising both elastic and inelastic scattering. A fundamental limitation of cryo-EM is the fact that inelastic scattering deposits energy in the specimen, causing irreversible sample damage and rapid deterioration of high-resolution details, a process termed radiation damage (R. M. Glaeser 1971; Henderson 1995). The deleterious effects of radiation damage on the imaged sample have been well documented by measuring the reduction of intensities of diffraction spots in exposure series obtained from 2D and thin 3D crystals (Unwin and Henderson 1975; Hayward and Glaeser 1979; Stark, Zemlin, and Boettcher 1996; Baker et al. 2010), as well as more recent measurements in noncrystalline, biological material (Grant and Grigorieff 2015).

Although radiation damage is unavoidable in cryo-EM imaging, its impact changes with different acceleration voltages. This is due to the fact that, although both elastic and inelastic scattering cross sections increase as a function of decreasing electron energy, the inelastic cross-section scales more slowly than the elastic cross-section (Peet, Henderson, and Russo 2019). In other words, there are more signal-producing elastic scattering events than damage/noise-producing inelastic scattering events at lower electron energies such as 100 keV, as compared to 300 keV, the energy most commonly used for high-resolution imaging. This relationship is well-established and has been observed through electron energy-loss spectroscopy (Reimer and Ross-Messemer 1989; 1990; Crozier 1990; Egerton 2011). More recent quantitative measurements of radiation damage and its dependence on acceleration voltages has determined that the elastic cross-section is 2.01-fold greater at 100 keV than at 300 keV, whereas the inelastic cross-section change is only 1.57-fold (Peet, Henderson, and Russo 2019). This implies that the amount of useful information from cryo-EM images of biological structures should be 25% greater using 100 keV compared to 300 keV electrons. Based on these findings, cryo-EM imaging using lower energies should theoretically yield more information per unit damage, yielding higher contrast in images that could enable more precise particle alignment of smaller molecular weight samples, something that is a current limitation of cryo-EM.

Although there are clear benefits of imaging biological samples at 100 keV, current direct detectors, which were crucial for the widespread adoption of cryo-EM were optimized for imaging at 300keV rather than at 100 keV. Therefore, current direct electron detectors perform poorly at 100 keV due to increased backscatter and side-scatter in the chip, leading to the detection of electrons in neighboring pixels and greatly reducing the detective quantum efficiency (DQE) at 100 keV (Peet, Henderson, and Russo 2019, Lee et al. 2022). Recent work in cryo-EM imaging at 100 keV has relied on repurposed hybrid-pixel X-ray detectors, but these hybrid-pixel arrays suffer from a small pixel area of only 0.5 megapixels (Mpix) (Naydenova et al. 2019). This smaller imageable area leads to a significant decrease in throughput, as well as less accurate CTF fitting due to suboptimal power in the amplitude spectrum. Furthermore, these customized hybrid-pixel arrays designed for applications in physics and materials science are suitable for practical demonstrations but are prohibitively costly for widespread use in cryo-EM. The answer to realizing the widespread use of cryo-EM imaging at 100 keV, to realistically capture the advantages of lower electron energies, is a commercially available, high-DQE, high-frame rate, high-area (>4 Mpix) detector.

Another major consideration in detector design is the need to drive down the detector cost. Since many institutions around the globe have invested in high-end instrumentation and there are numerous centralized service centers that provide access to high-end cryo-EM instrumentation, most researchers interested in pursuing single-particle cryo-EM studies are able to obtain datasets for image analysis. However, it is common knowledge in the field that the vast majority of samples require substantial optimization for effective data collection that will yield a high-resolution reconstruction (Carragher et al. 2019). Unless a research group has routine access to a local electron microscope with a high-end detector, this optimization process can severely delay progression of a cryo-EM project, particularly for smaller (< 200 kDa) complexes. Higher-end microscopes remain particularly expensive to purchase and maintain, in part due to the sophisticated autoloading device and associated autogrids required for loading cryo-EM samples for imaging. However, lower-end microscopes that use traditional side-entry cryo holders for grid insertion are maintained by many institutes that do not have the funding to support a higher-end microscope. A 100 keV microscope that uses a side-entry cryo holder could serve as an excellent tool if paired with a high-quality detector, enabling local cryo-EM sample optimization that confirms the high likelihood of a high-resolution structure when imaged on a more stable, higher-end instrument. Local screening in this manner would also substantially increase the efficacy of data collection on higher-end instruments.

Here, we characterize the recently developed Gatan Alpine direct detector (Gatan Inc., Pleasanton, CA) which has been specifically optimized to operate at 100-200 keV and be cheaper than the Gatan K3 detector. This optimization results in a 4-fold increase in DQE at the Nyquist frequency at 100 keV, as compared to the Gatan K3 which was optimized for 300 keV (Lee et al. 2022). Furthermore, this detector includes a large-format 7 Mpix sensor, much more suitable for the imaging throughput and accuracy of cryo-EM imaging. Excitingly, similarly optimized detector systems for cryo-EM at 100 keV (Dectris Singla with 1 Mpix) have already been shown to produce high-quality cryo-EM reconstructions (McMullan et al. 2023).

In this work, we have collected 11 datasets of a series of standard test samples and a non-standard biological sample closer approximating the challenging targets of the structural biology community, across a series of six microscope, detector, and accelerating voltage configurations, in order to evaluate the Gatan Alpine detector against current detector systems. We were able to obtain better than 2 Å resolution reconstructions of apoferritin collected with Gatan Alpine equipped on a ThermoFisher Scientific (TFS) Glacios microscope at both 120 and 200 keV, demonstrating a high level of performance of the detector. Additionally, we obtained 3.2 Å resolution reconstructions of a 115 kDa asymmetric protein complex collected using Gatan Alpine on a TFS Glacios at these voltages, demonstrating the applicability of this detector to more challenging biological samples. Remarkably, in-depth analysis of the data showed that collection with Gatan Alpine at 120 keV on a TFS Glacios was bested only by a collection at 300 keV using a Gatan K3 camera on a TFS Titan Krios with a BioQuantum Gatan energy filter, outperforming 200 keV TFS Glacios datasets collected with either Gatan Alpine or Gatan K3. To further demonstrate the utility of the Alpine detector on lower-end microscopes that are not equipped with expensive autoloading devices, we also determined a 2.6 Å resolution reconstruction of apoferritin and a 3.2 Å resolution reconstruction of aldolase on a TFS Talos F200C using a Gatan 626 side-entry holder. Our results demonstrate the applicability of the Alpine detector to structural biology on higher-end and lower-end systems.

## 2. METHODS

### 2.1. PROTEIN EXPRESSION AND PURIFICATION

#### 2.1.1 Murine heavy chain apoferritin for the Alpine/Glacios, K3/Glacios, K3/Titan Krios setups

Murine heavy chain apoferritin plasmid in pET24a vector was gifted from the laboratory of Radostin Danev and purified according to their published protocol (Danev, Yanagisawa, and Kikkawa 2019). Expression was carried out in E. coli BL21(DE3) cells as in the published protocol, and the final concentration step was carried out in a 50 kDa MWCO spin column and resulted in a ~10 mg/mL solution of apoferritin in 20 mM HEPES pH 7.5, 300 mM NaCl. This sample was then flash frozen in liquid nitrogen and stored at −80 ºC.

#### 2.1.2 LKB1-STRADα-MO25α for the Alpine/Glacios, K3/Glacios, K3/Titan Krios setups

A single plasmid with MO25α-P2A-6xHis-STRADα-P2A-FLAG-LKB1 (all full length) was transfected into 1 L Expi293T cell culture (Life Technologies). The cells were harvested and centrifuged at 1000 xg for 10min at 4 ºC, resuspended in ice-cold PBS and centrifuged again at 2000 xg for 10 min at 4 ºC. LKB1, STRADα, and MO25α proteins were then purified based on a previously published protocol (Zeqiraj et al. 2009). Briefly, cell pellet was resuspended in 150 mL of lysis buffer (50 mM Tris HCl pH 7.8, 150 mM NaCl, 5% glycerol, 1 mM EDTA, 1 mM EGTA, 10 mM BME, 20 mM imidazole) supplemented with 3 protease inhibitors (Roche) and then lysed by an EmulsiFlex-C3 (Avestin) at ~15,000 psi, 4 °C. The lysis was clarified through centrifugation at 35,000 xg for 30 min at 4 ºC. The supernatant was incubated rotating for 1 h at 4 ºC on 1.25 mL bed of HisPur Ni-NTA agarose beads (ThermoFisher Scientific), pre-equilibrated in lysis buffer. The beads were washed with 10 column volumes (CV) of a low salt buffer (50 mM Tris HCl pH 7.8, 150 mM NaCl, 5% glycerol, 1 mM EDTA, 1 mM EGTA, 10 mM BME, 20 mM imidazole) and 50 CV of a high salt buffer (low salt buffer containing 500 mM NaCl), followed by 10 CV of low salt buffer. The protein was eluted from the column with 12 mL of elution buffer (low salt buffer with 250 mM imidazole). The sample was diluted 2X and loaded onto a Mono Q 10/100 GL column (Cytiva). Bound proteins were eluted over 20 CV using a salt gradient of 0-700 mM NaCl and the LKB1-STRADα-MO25α complex eluted at ~200 mM NaCl. Peak fractions were pooled and concentrated to 500 µL and loaded onto a Superdex 200 Increase 10/300 GL (Cytiva), pre-equilibrated in 25 mM Tris-HCl pH 7.6, 350 mM NaCl, 5% glycerol, and 0.5 mM DTT. The LKB1-STRADα-MO25α complex eluted as a single peak. The peak fractions were pooled and concentrated to 3.68 mg/mL as determined by nanodrop A280 readings. The sample was aliquoted and flash frozen to be stored at −80 ºC.

#### 2.1.3 Murine heavy chain apoferritin for the Alpine/Talos setup

Murine heavy chain apoferritin plasmid in pET24a vector was received from Masahide Kikkawa (University of Tokyo), expressed and purified using a previously published protocol (Danev, Yanagisawa, and Kikkawa 2019). Vector was transformed into and expressed in BL21(DE3)pLysS *E. coli* chemically-competent cells according to the published protocol. Sample concentrated to 10-20 mg/mL was loaded onto a Superdex 200 Increase 10/300 (Cytiva) column equilibrated with 30 mM HEPES pH 7.5, 150 mM NaCl, 1 mM DTT. The fractions corresponding to the apoferritin peak were pooled and concentrated to 5 mg/mL. For long-term storage, trehalose (5% (v/v) final) was added to concentrated apoferritin prior to aliquoting and flash-freezing in liquid nitrogen for long-term storage at −70 ºC.

#### 2.1.4 Aldolase for the Alpine/Talos setup

Rabbit muscle aldolase was prepared as described previously (Herzik, Wu, and Lander 2017). Lyophilized rabbit muscle aldolase was purchased (Sigma Aldrich) and solubilized in 20 mM HEPES (pH 7.5), 50 mM NaCl at ~3 mg/ml. Aldolase was loaded onto a Sepharose 6 10/300 (Cytiva) column equilibrated in the solubilization buffer, and fractions corresponding to aldolase were pooled and concentrated to 1.6 mg/ml. Sample was aliquoted and kept on ice for immediate cryo-EM grid preparation.

#### 2.1.5 Transthyretin for the Alpine/Talos setup

Transthyretin in pMMHa vector was transformed into BL21(DE3) *E. coli* chemically-competent cells (New England BioLabs). Single colonies of transformed cells from agar plates containing 100 μg/mL ampicillin and grown at 37 °C were used for starter cultures using 50 mL LB media in 250 mL shake flasks containing 100 μg/mL ampicillin. The flasks containing the cultures were shaken at 37 °C until the media appeared turbid at which point 1 L shake flask containing 100 μg/mL ampicillin were inoculated with a 1:20 dilution of the starter culture and grown at 37 °C until OD600 was approximately 0.5. Cultures were induced with 1mM IPTG, incubated and shaken overnight at 30 °C, and harvested by centrifugation at 6,000 rpm for 30 minutes at 4 °C. The supernatant was removed, and the pellet resuspended with 50 mL TBS with an added protease inhibitor tablet (ThermoFisher Scientific Pierce Protease Inhibitor Tablets EDTA-Free). The resuspended pellet was sonicated three times with a Qsonica Q125, which involved 3 minutes of sonication followed by 3 minutes of rest at 4 °C. The sonicated sample was centrifuged at 15,000 rpm for 30 min and the supernatant collected. The supernatant was subjected to 50% ammonium sulfate precipitation (w/v) and then stirred for 45 minutes at room temperature. The sample was centrifuged at 15,000 rpm for 30 minutes at 4 °C, the supernatant collected and precipitated again with 90% ammonium sulfate (w/v), and then stirred for 45 minutes at room temperature. The sample was centrifuged, and the pellet collected and dialyzed with 3,500 MWCO dialysis tubing overnight in a buffer of 25 mM Tris pH 8.0 in a 4 °C cold room. The dialyzed sample was filtered with a low protein binding filter (Millipore Sigma) and run through a SourceQ15 anion exchange column (Cytiva). The sample was injected using a 5% Buffer A (25mM Tris pH 8.0) and eluted using a 60 min gradient up to 100% Buffer B (25 mM Tris/1.0 M NaCl) at room temperature. Eluates were then run through a Superdex 75 gel filtration column that had been equilibrated with 10 mM sodium phosphate pH 7.6 100 mM KCl. Peak fractions were collected for subsequent steps. A stabilizing stilbene substructure was covalently linked to Lys15 in each of the two binding pockets to stabilize the transthyretin tetramer. Ligand attachment involved adding 1 µL of a 1.5 mM solution of stilbene ligand in DMSO to 19 µL of purified transthyretin at a concentration of 1.7 mg/mL and incubating for 4 hours in the dark at room temperature. Transthyretin sample was aliquoted, flash frozen in liquid nitrogen, and stored at −70 ºC until thawed for cryo-EM sample preparation.

### 2.2 CRYO-EM GRID PREPARATION

#### 2.2.1 Apoferritin for the Alpine/Glacios, K3/Glacios, K3/Titan Krios setups

2 µL of purified murine apoferritin at a concentration of 8 mg/mL (in 20 mM HEPES 7.5, 300 mM NaCl) was applied to UltrAuFoil R 1.2/1.3 400 mesh (Alpine) or Quantifoil R 1.2/1.3 300 mesh (K3) Au holey-carbon grids (glow discharged for 30s at ~15mA before sample application) and blotted with Whatman No. 1 filter paper for ~6 seconds with blot force 0 on a Vitrobot Mark IV (ThermoFisher Scientific) at 100% humidity and 22°C.

#### 2.2.2 LKB1-STRADα-MO25α for the Alpine/Glacios, K3/Glacios, K3/Titan Krios setups

3 μL of purified heterotrimer complex at a concentration of 3 μM was applied to Quantifoil R1.2/1.3 400 mesh Au holey-carbon grids, blotted using a Vitrobot Mark IV (ThermoFisher Scientific) and plunge frozen in liquid ethane (15 mA 30 s glow discharge, 0 s wait time, 4 °C, 100% humidity, 6–8 s blot time, 4 blot force).

#### 2.2.3 Apoferritin and aldolase for the Alpine/Talos F200C setup

The apoferritin and aldolase samples imaged on the Talos F200C microscope were prepared on UltrAuFoil holey carbon grids (300 mesh, 1.2/1.3 µm spacing) purchased from Quantifoil. Grids were plasma cleaned for six seconds at 15 Watts (75% nitrogen/25% oxygen atmosphere) using a Solarus plasma cleaner (Gatan, Inc.) immediately prior to sample preparation, which was performed using a custom-built manual plunger located in a cold room (≥95% relative humidity, 4 °C). 3 µL of purified aldolase (1.6 mg/mL) or apoferritin (5 mg/mL) was applied to grids and manually blotted for four to five seconds using Whatman No. 1 filter paper. Blot time counting commenced once the blotted sample on the filter paper stopped spreading, monitored visually using a lamp positioned behind the sample. At the end of 4-5 s, the blotting paper was pulled back and immediately plunged into a well of liquid ethane cooled by liquid nitrogen.

#### 2.2.4 Transthyretin for the Alpine/Talos F200C setup

TTR tetramers exhibit a preferred orientation due to interactions with the air-water interface when prepared on traditional holey carbon grids, so the TTR sample was frozen on R1.2/1.3 UltrAuFoil Holey Gold grids (Quantifoil) that had been coated with a single layer of monolayer graphene. Graphene grids were prepared as described previously (Basanta et al. 2023). The monolayer graphene was partially oxidized prior to cryo-EM sample preparation by exposure to UV/ozone for four minutes using a UVOCS T10×10 ozone generator. TTR cryo-EM grids were manually plunged in a cold room as for the 100 keV apoferritin and aldolase samples. The grids were blotted for 6 seconds using a 1 × 6 cm piece of Whatman No. 1 filter paper and plunged in liquid ethane.

### 2.3 CRYO-EM DATA COLLECTION (see Supplemental Table S1)

For all datasets collected with the Gatan Alpine detector on the Glacios microscope (equipped with a narrow pole piece with the gap of 7 mm, called the SP-Twin lens) at 120 keV or 200 keV, the C2 aperture was 70 μm and the objective aperture was 100 μm. Comparison datasets were collected with the Gatan K3 detector on the Glacios microscope at 200 keV or the Titan Krios G2 with a BioQuantum Imaging Filter and energy filter slit width of 20 eV. With the K3 Glacios setup, the C2 aperture was set to 30 μm and the objective aperture size was 100 μm. With the Titan Krios setup, the C2 aperture was 70 μm and the objective aperture size was 100 μm. Data was collected automatically using the 3×3 9-hole image shift in SerialEM *v3*.*8*.*6* with beam tilt compensation (Mastronarde 2005).

For all datasets collected with the Gatan Alpine detector on the Talos F200C microscope at 100 keV or 200 keV, the C2 aperture was 50 µm, and the objective aperture was 100 µm. Data was collected automatically using the Leginon data acquisition software (Cheng et al. 2021) using 5-hole image shift with beam tilt compensation to counter the introduction of image shift-dependent coma. Representation of data collection can be seen in Supplemental Table 1.

#### 2.3.1 Edge-method DQE measurements

The DQE of the Alpine and K3 cameras was measured in super-resolution counting correlated-double sampling (CDS) mode, using knife-edge images. The DQE measurements were performed using the method described in (Mooney 2007) and ISO 12233 (https://www.iso.org/obp/ui/#iso:std:iso:12233:ed-1:v1:en). The edge-method DQE is known to slightly exaggerate the DQE at zero spatial frequency. To account for the non-linear response of electron counting cameras, a dose rate within the linear regime of the camera was selected. All measurements were conducted at a dose rate of 7.5 electrons pixel^−1^ second^−1^ to maintain uniform calibration across the evaluations.

#### 2.3.2 Microscope alignment

Full alignments of microscopes were performed at 100 kV, 200 kV (Talos F200C); 120 kV, 200 kV (Glacios); and 300 kV (Titan Krios). Parallel illumination was used for all datasets collected, either by setting the C2 current in nanoprobe on the two-condenser Glacios or Talos F200C as published (Herzik, Wu, and Lander 2017) or using the parallel range on the three-condenser Krios. For alignments immediately before data collection on the Glacios and Krios, the sample was first set to a stage position at eucentric height over carbon, specimen was then brought close to focus and pivot points and rotation center were adjusted at imaging magnification using a beam intensity that corresponded to parallel illumination. The SerialEM implementation of astigmatism and Coma-free alignment correction by CTF fitting was used for fine adjustment, correcting 8 beam tilt directions. SerialEM Coma vs. Image Shift procedure was then performed. For alignments immediately prior to data collection on the Talos F200C, a cross-line grating replica grid with Au/Pd shadowing was inserted with a room temperature side-entry holder and brought to eucentric height. Pivot points and rotation center were adjusted at focus, and parallel illumination was set in diffraction mode using the objective aperture and gold diffraction. Coma-free alignment was performed using a Zemlin tableau with the Leginon software as previously described (Robert M. Glaeser et al. 2011).

#### 2.3.3 Consideration for data collection consistency

For consistency, apoferritin Glacios and Krios datasets were collected from the same sample purification at UCSF, and the apoferritin Talos F200C datasets were collected from the same sample purification at Scripps. Alpine grids were frozen during a single session at each site; K3 grids were frozen in a separate session.

For consistency, LKB1 complex datasets were collected from the same sample purification and grids were vitrified under the same conditions in two freezing sessions at UCSF.

### 2.4 IMAGE PROCESSING (see Supplemental Table S2, S3, S4)

During data collection on the Glacios and Krios, the image quality was estimated in real time using Scipion software (de la Rosa-Trevín et al. 2016). Image frames acquired on the Talos F200C were preprocessed and monitored for data quality in real-time using the Appion software (Lander et al. 2009). Beam-induced motion correction and dose weighting were performed on the raw movie stacks on the fly using MotionCor2 (v1.4.0) (Zheng et al. 2017). For collection with the K3 camera, a patch size of 7×5 with no grouping of frames was used. Since the sensor for the Alpine is smaller and has differing dimensions, we used a 3×4 patch size with a grouping of 3 frames. All datasets had a b-factor of 100 with 7 iterations. Whole image contrast transfer function (CTF) estimation was performed using CTFFIND4 (Rohou and Grigorieff 2015). Prior to initial single particle data collection, rough pixel sizes for each magnification were calculated based on gold diffraction from a cross-line grating with Au shadowing. Following collection of Apoferritin datasets, pixel sizes were determined more precisely by cross correlation with a published high-resolution Apoferritin dataset, EMDB-11668 (Yip et al. 2020), and our experimental Apoferritin maps using the “Fit in Map” operation in Chimera.

#### 2.4.1 Apoferritin data collection and image processing on Alpine/Glacios, K3/Glacios, K3/Titan Krios setups

Apoferritin data was collected in super resolution CDS mode for the Alpine and non-CDS mode for the K3 data sets. All image processing after motion correction was performed with cryoSPARC v4.2 (Punjani et al. 2017). PatchCTF was used to perform CTF estimation. Exposures with fits below 3.5 Å (Krios) or 6 Å (Glacios) were removed, leaving 3312 micrographs for Alpine 120 keV, 3619 for Alpine 200 keV, 3903 for Glacios/K3 200 keV, and 617 for Krios. Particles were picked with Blob picker (1,635,971, Alpine 120; 1,834,404, Alpine 200; 2,675,691, Glacios K3; 790,751, Krios K3) and subjected to 2 rounds of 2D classification, leaving 302,338, Alpine 120; 204,387, Alpine 200; 686,940, Glacios K3; 245,959, Krios K3. 50,000 particles were used to create an *ab initio* model with O symmetry, which was then used as a reference map for symmetric homogeneous refinement with both global CTF refinement (fit tilt, trefoil, spherical aberration, tetrafoil, and anisotropic magnification) and per-particle defocus refinement enabled. Local CTF refinement was repeated, and these particles were fed into a final homogeneous refinement job with all previous settings and Ewald sphere correction turned on (curvature sign was determined with two reconstruction jobs with each curvature sign; the one with the highest resolution was used for future refinement). To remove bias from differing particle number for the final reconstructions, all datasets were trimmed down to ~200,000 particles and homogeneous refinement was repeated with the same settings.

#### 2.4.2 LKB1 data collection and image processing on Alpine/Glacios, K3/Glacios, K3/Titan Krios setups

LKB1-STRADα-MO25α data for the Alpine datasets was collected in super resolution CDS mode. For Alpine 120 keV dataset, image processing was performed in cryoSPARC v4.2 (Punjani et al. 2017) software following the standard protocols. Patch-based CTF estimation was completed in cryoSPARC using PatchCTF. Micrographs were bulk curated with a CTF fit better than 8 Å and manually curated based on particle presence and distribution to yield 2,868 micrographs. 897,749 particles were picked using a template picker with templates of 2D classes from a previous LKB1 dataset that were low pass filtered to 30 Å and extracted at 2x binned. These particles were 3D classified to eliminate “junk” classes and particles through iterative rounds of ab initio reconstruction and heterogenous refinement. Particles associated with the selected good class were reextracted unbinned with a box size of 256 pixels with further rounds of *ab initio* reconstruction and heterogenous refinement. Non-uniform refinement was used for the final reconstruction that yielded a final average resolution of 3.19 Å (FSC= 0.143) from 120,684 particles.

For data processing for the Alpine 200 keV dataset (collected in CDS mode), and K3 200 keV and K3 300 keV datasets (collected in non-CDS mode), the images were processed in cryoSPARC following the same protocol as above. Micrographs were bulk curated with a CTF fit better than 8 Å and the number of manually curated micrographs (3,179 Alpine/Glacios 200 keV; 1,337 K3/Glacios 200 keV; 1,223 K3/Krios) used for the reconstructions were based on having similar initial particle stack sizes for all comparison datasets (1,229,097 Alpine/Glacios 200 keV; 1,000,313 K3/Glacios; 979,927 K3/Krios) (Supplemental Table 3). The same LKB1 templates were used for template picking in all datasets. Due to slight differences in angstrom/pixel sizes, slightly different box sizes were used but all between 217 Å and 221 Å. All final reconstructions were created from non-uniform refinement jobs in cryoSPARC and contained similar number of particles (120,684 Alpine 120 keV; 111,117 Alpine/Glacios 200 keV; 111,117 K3/Glacios; 124,780 K3/Krios) (Supplemental Table 3). All the reported resolutions are based on the 0.143 Fourier shell criterion (Chen et al. 2013; Rosenthal and Henderson 2003) (Supplemental Figures 2,3).

#### 2.4.3 Apoferritin data collection and image processing on the Talos F200C/Alpine setup at 200 keV

An apoferritin grid was loaded into a Gatan 626 side-entry cryo holder and inserted into a TFS Talos F200C microscope for data collection at 200 kV. For data collection, the liquid nitrogen in the cryo-holder dewar was maintained under the sample rod and refilled every 1.5 hours. Data collection was paused during manual refilling and restarted once the liquid nitrogen in the dewar had a glass-like surface and the drift rate had decreased to less than 1 Å/sec. Data was collected in super resolution CDS mode, at a nominal magnification of 45,000x (0.851 Å/pixel).

100 movies were acquired in super-resolution CDS mode at a nominal magnification of 45,000x (0.872 Å/pixel) using a defocus range −0.8 µm to −1.4 µm with an exposure rate of 7.81 e^−^/pixel/s for a total of 5 seconds (100 ms/frame, 50 frames), resulting in a total exposure of ~51 e^−^/Å^2^ (1.02 e^−^/Å^2^/frame). Frame alignment with dose-weighting was performed using the MotionCor2 frame alignment program (v1.3.1) (Zheng et al. 2017) with 3 × 4 tiled frames, a B-factor of 500, and without frame grouping, as implemented within Appion. The summed, dose-weighted images were then imported into cryoSPARC v.3.2.0 (Punjani et al. 2017) for Patch CTF estimation (0.07 amplitude contrast, 30 Å minimum resolution, 4 Å maximum resolution). Aligned images with a CTF maximum resolution worse than 4 Å were excluded from further processing.

2D class averages of apoferritin generated from a prior dataset acquired on an Arctica microscope with an identical pixel size were used for template picking in cryoSPARC using default parameters. A total of 29,404 particle selections were extracted from 84 micrographs, binned 2 × 2 (1.744 Å/pixel, 180-pixel box size), and subjected to reference-free 2D classification (50 classes). *Ab initio* models were generated from selected (21,174 particles) as well as rejected particles from 2D classification, and used as reference models for heterogeneous refinement (2 classes, C1 symmetry). The coordinates of the better-resolved class comprising 21,133 particles were then imported into RELION 3.1 (Zivanov et al. 2018) and re-extracted un-binned (360-pixel box size). 3D auto-refinement with O symmetry yielded a ~3.5 Å reconstruction. Per-particle defocus, beam tilt, and trefoil aberration estimation were subsequently performed in RELION. To more accurately measure beam tilt for each set of image shift targets, particles were sorted into 5 optics groups according to beam-image shift parameters used during acquisition (average residual beam tilts estimated as x=0.24 and y=-0.39 mrad). Particles with optimized optics corrections were imported into cryoSPARC and homogenous refinement with O symmetry yielded a ~2.7 Å reconstruction according to a gold-standard FSC at 0.143 (Supplemental Figure 4, Table S4). Performing tetrafoil and Ewald sphere correction during refinement did not have an impact on the reported resolution or quality of the reconstruction density. The local resolution ranged from 2.4 to 2.8 Å in the best and worst-resolved regions (Supplemental Figure 4), and the estimated B-factor, based on the slope of the Guinier plot from 7 to 2.7 Å, was 90.9.

#### 2.4.4 Apoferritin data collection and image processing on the Talos F200C/Alpine setup at 100 keV

A new apoferritin grid was loaded into a Gatan 626 side-entry cryo holder and inserted into a TFS Talos F200C microscope for data collection at 100 kV. Nitrogen level, refilling, and drift rate monitoring was performed as for the apoferritin. The pixel size of the Alpine images on the Talos F200C at 100 keV was determined following the steps used to determine the pixel size on the Glacios at 120 keV, using a high-resolution structure that was initially determined using the 200 keV pixel size. Data were then reprocessed using the same methodology using the correct pixel size. 318 movies were acquired in super resolution CDS mode at 45,000x (0.852 Å/pixel) at 100 keV using a defocus range −0.3 µm to −0.9 µm with an exposure rate of 5.2 e^−^/pixel/second, for a total of 20 sec (100 ms/frame, 200 frames), resulting in a cumulative dose of ~143 e^−^/Å^2^ (0.72 e^−^/Å^2^/frame). More frames than typically used for data collection were acquired to compare image analysis using different ranges of frames, but processing with frames contributing to a cumulative exposure of 51 e^−^/Å^2^ gave the same results as when all frames were used. As for the 200 keV apoferritin dataset, frame alignment with dose-weighting was performed on 3 × 4 tiled frames with a B-factor of 500 (no frame grouping) using the MotionCor2 v1.3.1 (Zheng et al. 2017). Since MotionCor2 v1.3.1 does not recognize 100 keV as a valid voltage input and defaults to applying the critical dose calculated for 300 keV for dose-weighting, we used a cumulative exposure of 1.57 times the total exposure as input. Since this is an imprecise workaround, it is possible that determining the dose curve for 100 keV would likely improve the final resolution of our processing.

The summed, dose-weighted images were then imported into CryoSPARC v.3.2.0 (Punjani et al. 2017) for Patch CTF estimation (0.1 amplitude contrast, 25 Å minimum resolution, 4 Å maximum resolution). Blob picker using a diameter of 110 Å was used to pick particles to generate ten 2D class averages for subsequent template picking. The first three 2D averages were used for template-based particle selection using the default parameters, and a total of 82,977 template picks were extracted un-binned from 218 micrographs (256-pixel box size), which were subjected to two rounds of 2D classification (50 classes each). *Ab initio* models were generated from the 59,000 particles contributing to detailed 2D averages, as well as rejected particles from 2D classification, and used as reference models for heterogeneous refinement (2 classes, octahedral symmetry imposed). Particles corresponding to the better-resolved class (51,253 particles, ~3.9 Å) were then subjected to homogeneous refinement with O symmetry. Per-particle defocus estimation, and per-group CTF refinement correcting for beam tilt, trefoil, spherical aberration (C_s_), and tetrafoil aberrations, yielding a ~2.6 Å reconstruction according to gold-standard FSC at 0.143 (average residual beam tilts estimated as x=0.32 and y=-0.44 mrad) (Table S4). Performing tetrafoil and Ewald sphere correction during refinement did not have an impact on the reported resolution or quality of the reconstruction density. The local resolution ranged from 2.4 to 2.8 Å in the best and worst-resolved regions (Figure 5A), and the estimated B-factor, based on the slope of the Guinier plot from 7 to 2.6 Å, was 103.4.

#### 2.4.5 Aldolase data collection and image processing on the Talos F200C/Alpine setup at 100 keV

An apoferritin grid was loaded into a Gatan 626 side-entry cryo holder and inserted into a TFS Talos F200C microscope that had been aligned at 100 kV. Nitrogen level, refilling, and drift rate monitoring was performed as for the apoferritin. 2,476 movies were collected in super resolution CDS mode at 45,000x (0.852 Å/pixel) using a defocus range −0.5 µm to −1.5 µm with an exposure rate of 5.2 e^−^/pixel/second. For each exposure, the sample was exposed for 10 sec (100 ms/frame, 100 frames), resulting in a cumulative dose of ~71 e^−^/Å^2^ (0.71 e^−^/Å^2^/frame). Motion correction was performed as for the 100 keV apoferritin dataset. Aligned, dose-weighted summed frames was imported into cryoSPARC v3.3.1 and CTF estimation was performed with CTFFIND4 (Rohou and Grigorieff 2015) using all default parameters.

Only the 2,027 micrographs with CTF fit values of 10 Å or better were retained for further processing. Particles were selected from the first 387 micrographs using the cryoSPARC blob picker using default parameters except the minimum and maximum diameters set to 70 Å and 100 Å, respectively, and the “use elliptical blob” option selected. Particles were extracted from these micrographs with a box size of 160 pixels and 2D classified into 50 classes with all default parameters. The 16 class averages that contained detailed secondary structural information were selected and used for template-based particle picking using a particle diameter of 100 Å (all other parameters default), resulting in 1,914,721 particle selections. The 966,964 particles with NCC scores greater than 0.4 and with “local power” values between 60 and 126 were extracted with a box size of 256 pixels. 2D classification of these extracted particles into 200 classes was performed with a window inner and outer radius of 0.7 and 0.8, respectively, and circular mask diameter set from 100 Å to 120 Å.

The particles contained in the 75 class averages containing detailed structural information were used to perform ab initio initial model generation, requesting three volumes with a window inner and outer radius of 0.7 and 0.85, respectively. Given that this sample was previously imaged for high-resolution reconstruction on other microscopes and was confirmed to have D2 symmetry, this symmetry was imposed during ab initio model generation. All other parameters were left with default values. One of the three resulting ab initio maps resembled the known structure of aldolase, while the other two appeared to be symmetrized non-particles. Heterogeneous refinement was next performed using four initial volumes. The first map was the ab initio model that most closely resembled aldolase, and the remaining three initial volumes comprised the non-particle reconstructions. Window radii of 0.7 and 0.85 were used, and D2 symmetry was enforced. 674,124 particles classified to the first volume, improving the resolution of the map to ~3.8 Å. These particles were used for non-uniform refinement with D2 symmetry imposed, and all other parameters left as defaults except “Do symmetry alignment” turn off and “initial lowpass resolution” set to 6 Å, yielding a 3.3 Å resolution structure. The stack was then imported into RELION 3.1 (Zivanov et al. 2018) and particles were sorted into 5 optics groups according to beam-image shift parameters stored in the Leginon database. Per-particle defocus, beam tilt, and trefoil aberration estimation were subsequently performed, and 3D auto-refinement with D2 symmetry and CTF refinement including 4^th^-order (symmetrical) aberration correction yielded a ~3.1 Å reconstruction according to gold-standard FSC (average residual beam tilts estimated as x=0.66 and y=-0.48 mrad) (Table S4). The local resolution ranged from 2.7 to 3.3 Å in the best and worst-resolved regions (Supplemental Figure 5B), and the estimated B-factor, based on the slope of the Guinier plot from 7 to 3 Å, was 189.2.

#### 2.4.6 Transthyretin data collection and image processing on the Talos F200C with the Alpine detector at 100 keV

A transthyretin grid was loaded into a Gatan 626 side-entry cryo holder, inserted into a TFS Talos F200C microscope aligned at 100 kV. Nitrogen level, refilling, and drift rate monitoring was performed as for the apoferritin and aldolase samples. 7,001 movies were collected in super resolution CDS mode at 45,000x (0.852 Å/pixel) using a defocus range −0.5 µm to −1.5 µm with an exposure rate of 5.2 e^−^/pixel/second. For each exposure, the sample was exposed for 7 sec (100 ms/frame, 100 frames), resulting in a cumulative dose of ~50 e^−^/Å^2^ (0.71 e^−^/Å^2^/frame).

RELION 3.1 was used to process the first 300 micrographs acquired to assess the quality of the data and attainable structural details as data was being acquired. Frames were imported into RELION and MotionCor2 1.3.1 was used for frame alignment and dose weighting, with 3 × 4 tiled frames, a B-factor of 500, and without frame grouping. Aligned, dose-weighted micrographs were imported into cryoSPARC v3.3.1 and CTF estimation performed with CTFFIND4 (0.1 amplitude contrast, min defocus 2000 Å, max defocus 20000 Å, all others default). Images with estimated CTF fits worse than 10 Å or defocus values lower than 0.5 µm were discarded. Particles were selected using the cryoSPARC blob picker with default parameters except the minimum and maximum diameters set to 50 Å and 80 Å, respectively. Particles were extracted from these micrographs with a box size of 160 pixels and 2D classified into 50 classes with all default parameters. The 8 class averages in which secondary structural features were visible were selected and used for template-based particle picking using a particle diameter of 70 Å (all other parameters default), resulting in 214,296 particle selections. The 142,390 particles with NCC scores greater than 0.49 and with “local power” values between 38 and 78 were extracted with a box size of 192 pixels. 2D classification of these extracted particles into 50 classes was performed with the following non-default settings: circular mask diameter set from 75 Å to 90 Å, force max over poses/shifts=off, number of online-EM iterations=80, batchsize per class=1000.

The full dataset of 7,001 movies was imported into RELION 3.1 and motion correction performed as for the first 300 movies. Aligned, dose-weighted micrographs were imported into cryoSPARC v3.3.1 and CTF estimation was performed as before. The 10 best class averages from the initial processing were used as templates for particle picking, yielding 2,944,250 particles. The 142,390 particles with NCC scores greater than 0.49 and with “local power” values between 38 and 78 were extracted with a box size of 192 pixels for further processing. However, a high-resolution structure was not obtained after extensive processing, including re-importing into Relion and performing Bayesian polishing.

### 2.5 ATOMIC MODEL REFINEMENTS

#### 2.5.1 Apoferritin

For apoferritin, the 1.1 Å resolution cryo-EM structure (PDB 7A6A) was rigid body fit into each cryo-EM density map in UCSF ChimeraX.

#### 2.5.2 LKB1-STRADα-MO25α

For LKB1 complex, the x-ray crystal structure PDB 2WTK was docked into the cryo-EM density map of the K3 Titan Krios 300 keV dataset in UCSF ChimeraX (Pettersen et al. 2004) and then flexibly fit in torsion space using Rosetta into the density with a weight of 50 (DiMaio et al. 2015). The model was further refined in Isolde 1.0 (Croll 2018) through a plugin for UCSF ChimeraX and additional residues were built into the map through Coot (Emsley and Cowtan 2004). Iterative rounds of Isolde, Rosetta FastRelax, and Realspace refine in COOT with manual examination were used to improve the fitting of the model while maintaining realistic geometries. The final structure was validated using phenix (Afonine et al. 2018). For each of the other dataset reconstructions, this model was further flexibly fit in torsion space using Rosetta into the density of each of the other maps for calculation of the Q-scores. Only the model associated with the highest resolution cryo-EM map was deposited into the PDB.

#### 2.6 QUANTITATIVE METRICS

For each dataset, the local resolution was estimated using the Local Resolution job in cryoSPARC (Punjani et al. 2017). Q-scores (Pintilie et al. 2020) were calculated for each map and model in Chimera (Pettersen et al. 2004). per-particle spectral signal to noise (ppSSNR) plots (Baldwin and Lyumkis 2020) were generated by dividing each reconstruction into subset particle stacks of around 30,000, 45,000, or 60,000 particles each, and calculating Fourier Shell Correlation (FSC) curves for each subset reconstruction using the ResLog Analysis job in cryoSPARC (Stagg et al. 2014). Noise-substituted FSC curves were converted into SSNR data (SSNR=FSC/1-FSC), and used to generate ppSSNR data by dividing the SSNR data by the number of particles for each subset reconstruction. ppSSNR data were averaged across subset reconstructions for each dataset, as the ppSSNR is independent of the number of particles in the subset reconstruction. Error bars were generated using standard deviation of the average ppSSNR for each dataset. These data were then plotted as the ln(ppSSNR) by both spatial frequency and fractions of physical Nyquist based on the nominal physical pixel sizes for each collection.

### 2.7 DATA AND CODE AVAILABILITY

Three-dimensional cryo-EM density maps for the LKB1 complex were deposited in the Electron Microscopy Data Bank (EMDB) under the accession numbers EMD-43506 (K3/Krios), EMD-43702 (K3/Glacios), EMD-43701 (Alpine/200 keV/Glacios) and EMD-43700 (Alpine/120 keV/Glacios). Atomic coordinates for the atomic model were deposited in the RCSB Protein Data Bank under the accession number PDB 8VSU (K3/Krios). Three-dimensional cryo-EM density maps for the apoferritin collected at UCSF were deposited in the Electron Microscopy Data Bank (EMDB) under the accession numbers EMD-43576 (Alpine/Glacios at 120 keV), EMD-43685 (Alpine/Glacios at 200 keV), EMD-43688 (K3/Glacios at 200 keV) and EMD-43703 (K3/300 keV/Krios). The apoferritin and aldolase maps acquired at 100 keV were deposited to the EMDB with the accession numbers EMD-43527 and EMD-43528, respectively. Any additional information required to reanalyze the data reported in this paper is available from the lead contact upon request.

## 3. RESULTS

### 3.1 Detective Quantum Efficiency (DQE) of the Gatan Alpine direct detection camera

The Gatan Alpine direct detection camera was developed to enable cryo-EM imaging within the 100-200 keV range. We determined the DQE of the Alpine camera at both 100 and 200 keV in super-resolution CDS mode (Fig. 1). The DQE of the Alpine at 200 keV is slightly higher at high spatial frequencies than at 100 keV. Interestingly, at low spatial frequencies, the DQE of Alpine is marginally higher at 100 keV. Since the Alpine sensor was optimized for 100-200 keV applications, it outperforms the K3 camera at 100 keV at all spatial frequencies, facilitating high-resolution structure determination at lower electron voltages.

**Fig. 1.**
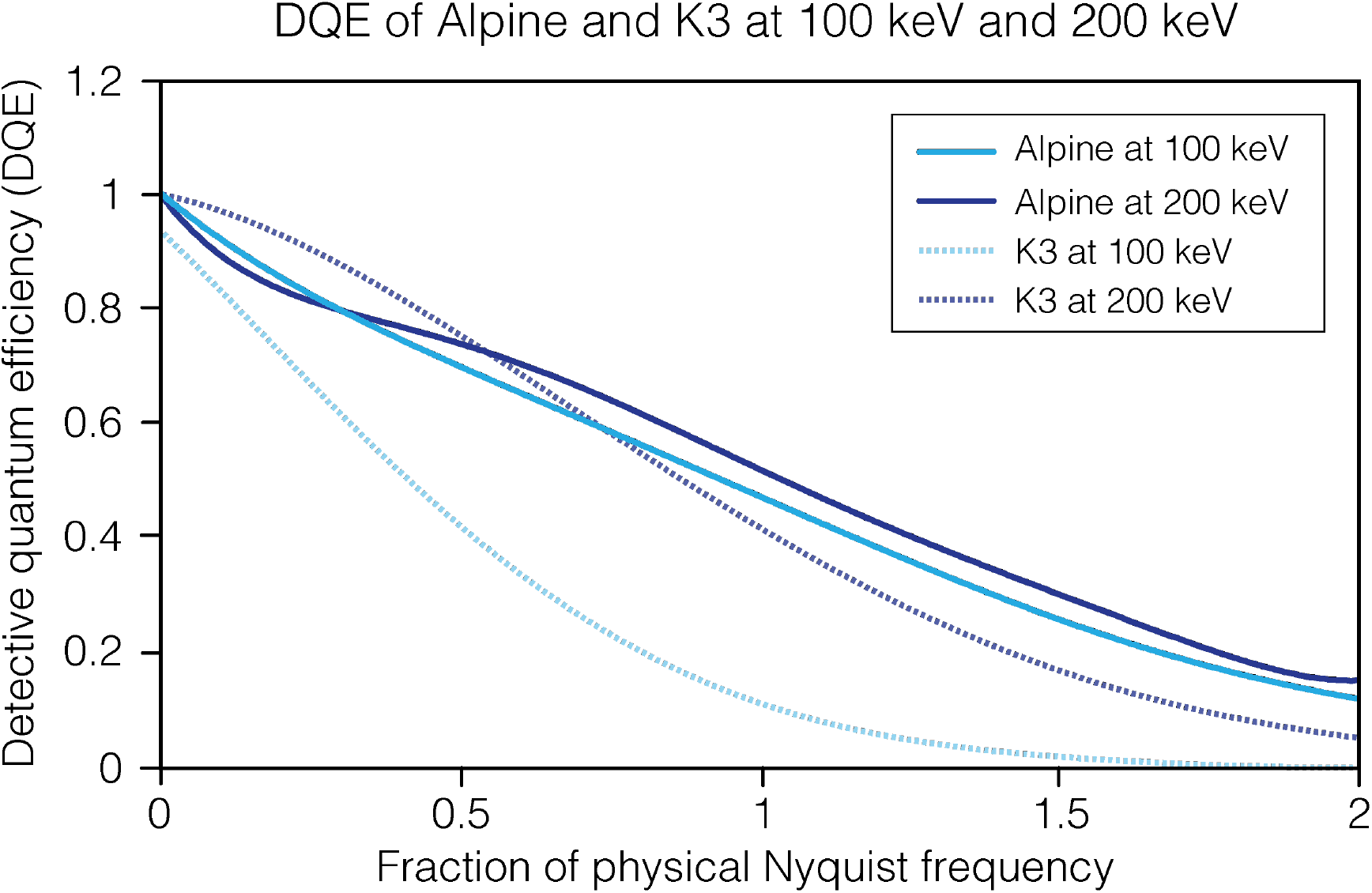
Detective Quantum Efficiency (DQE) of the Alpine and K3 camera. DQEs at 100 and 200 keV for Gatan Alpine direct detector and Gatan K3 direct detector across different fractions of physical Nyquist. DQEs were measured in super-resolution counting CDS mode at 7.5 electrons pixel^−1^ second^−1^.

### 3.2 Apoferritin collected with Gatan Alpine at 120 keV and 200 keV

To test the practical application of lower accelerating voltage electron detection by the Gatan Alpine sensor, we imaged apoferritin with the Alpine direct detector on a TFS Glacios microscope equipped with a narrow pole piece set to an accelerating voltage of 120 kV. We were able to obtain a high-resolution 3D reconstruction from 200,000 particles (trimmed to be consistent across data sets), with an estimated resolution of 1.92 Å (Fig. 2A, Supplemental Fig. 1A). The local resolution of the map ranges between 1.2 Å and 4.7 Å, with the majority of the map being between 1.5-1.9 Å. The Q-score of 0.85 is a quantitative measure confirming the level of structural resolvability expected at this resolution. We additionally imaged apoferritin with the Alpine detector on a TFS Glacios at 200 keV, which reached a resolution of 1.76 Å with 204,387 particles, (Fig. 2B, Supplemental Fig. 1B). The Q-score of 0.88 matches an estimated resolution of 1.35 Å, and the majority of the estimated local resolution (which ranged from 1.15 Å to 5.49 Å) was 1.5 to 1.6 Å. We note that Ewald sphere correction improved the resolution of both Alpine datasets between 0.05 Å (120 keV) and 0.07 Å (200 keV). It is important to note that these samples were collected on UltrAuFoil grids, which are known to decrease beam-induced motion (Russo and Passmore 2014) and likely contributed to increased image quality as compared to apoferritin data sets collected with the Gatan K3 detector (see comparisons below).

**Fig. 2.**
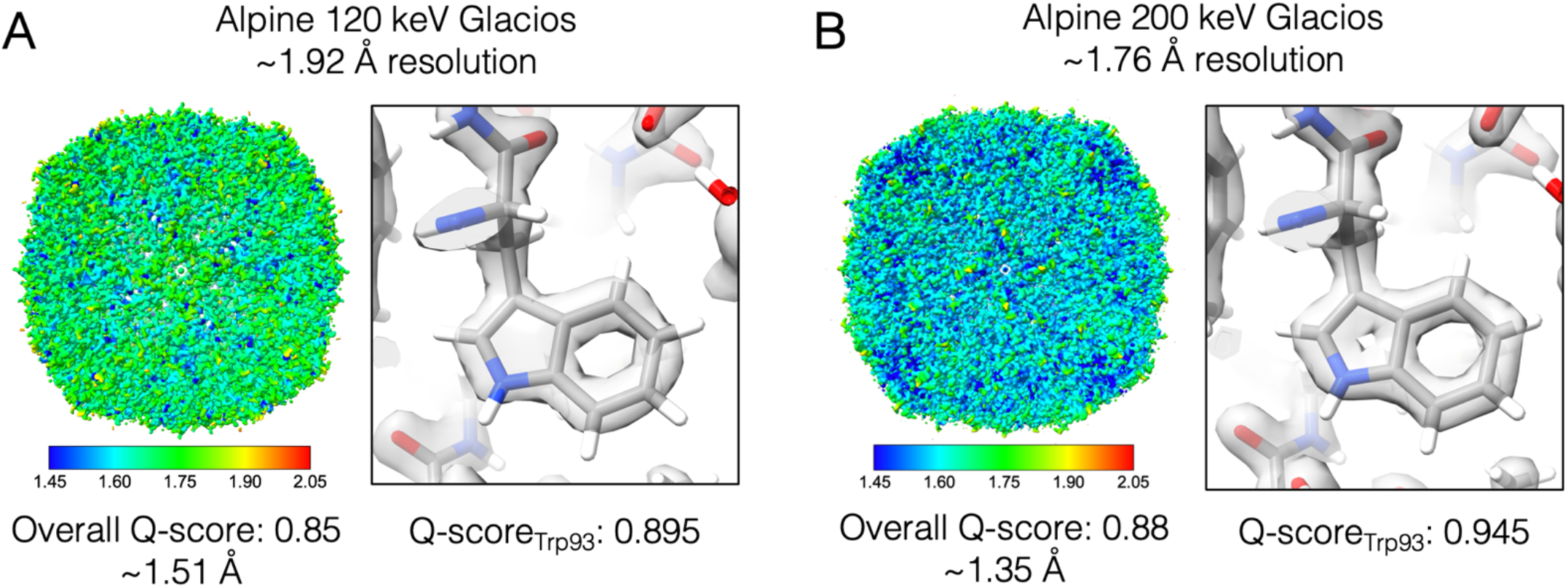
High-resolution reconstructions of apoferritin from Alpine detector on Glacios microscope at 120 and 200 keV. 3D reconstructions of apoferritin sample colored by the resulting local resolution from (A) 120 keV collection and (B) 200 keV collection. The average Q-score and the resolution associated with that Q-score is reported below each reconstruction. Close-up view of residue Trp93 fit into each density map shows high resolution features.

### 3.3 Comparison of LKB1-STRADα-MO25α complex reconstructions across different microscope and detector configurations

In addition to the collection with the standard apoferritin sample, we also tested the performance of the Gatan Alpine detector on the LKB1-STRADα-MO25α complex (referred to hereafter as the LKB1 complex), which is considerably smaller (115kDa) and asymmetric. With identical microscope collection parameters as the apoferritin datasets, we imaged the LKB1 complex with the Alpine detector on a TFS Glacios microscope at 120 keV and 200 keV. We collected the same number of micrographs for both collections (3,635) and curated micrographs to start with comparable initial particle amounts. Similar data processing workflows in cryoSPARC amounted to final particle stacks of similar sizes, with 120,684 particles for the 120 keV dataset and 111,117 particles for the 200 keV dataset (Supplemental Table 1). Similar to the apoferritin dataset, we obtained high-resolution reconstructions at both accelerating voltages. Notably, the LKB1 sample was resolved to a higher resolution at 120 keV than at 200 keV (3.2 Å vs. 3.4 Å, respectively), with the local resolution ranges and Q-score metrics following similar trends as the FSC resolutions (Fig. 3A,B, Supplemental Fig. 2).

**Fig. 3.**
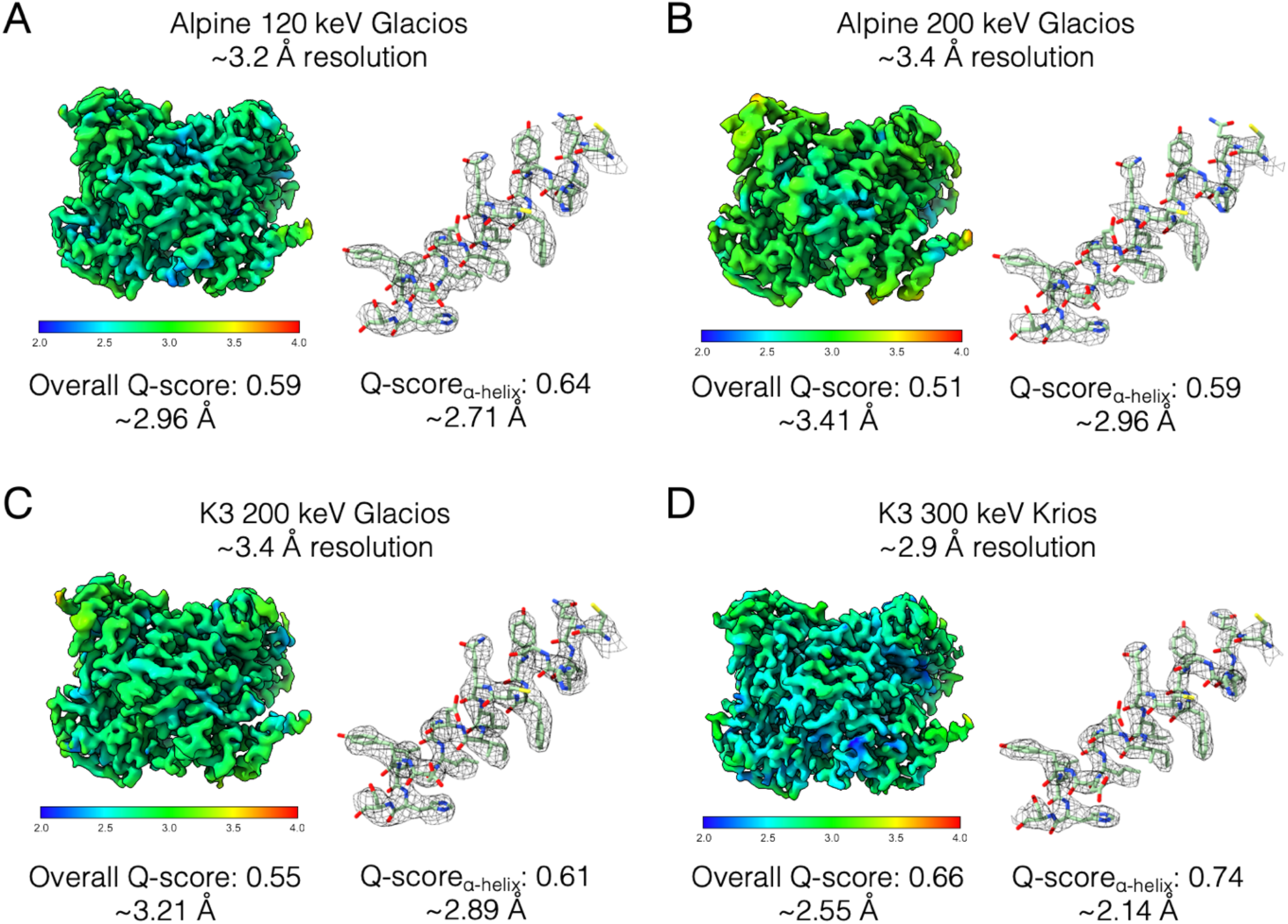
Reconstructions of LKB1 complex across multiple microscope/detector configurations at a range of accelerating voltages. 3D reconstructions of LKB1 heterotrimeric complex are colored by local resolution. To the right of each structure is a close-up view for a representative α-helix fit into each density map. Overall Q-scores and Q-score derived effective resolution are shown below each structure and the representative helix. In (A) and (B) are reconstructions from TFS Glacios microscope with the Alpine detector at 120 keV and 200 keV respectively. In (C) and (D) are reconstructions from the K3 detector collections on TFS Glacios at 200 keV and TFS Titan Krios at 300 keV respectively.

To compare the Alpine detector to current detectors, we collected additional datasets of the LKB1 complex using the K3 direct detector on a TFS Glacios at 200 keV. As the K3 detector is ~3 times the size of the Alpine detector, we collected a smaller dataset of 1,504 micrographs to have a comparable sized initial particles stack (Supplemental Table 1). 2D and 3D classifications resulted in comparable final particle stacks. The K3 200 keV Glacios reconstruction reached 3.37 Å, the same resolution as the Alpine 200 keV Glacios dataset (Fig. 3C). However, based on the local resolution estimates, we observed that the K3 cryo-EM map was of higher quality in most areas. We can quantitatively see this trend in the higher Q-score of 0.55 for the K3 collection compared to 0.51 for the Alpine collection at 200 keV (Fig. 3B,C, Supplemental Fig. 3A). Based on these results, the K3 outperformed the Alpine at 200 keV for this sample. However, the Alpine at 120 keV still yielded the highest resolution reconstruction as compared to either the K3 or Alpine 200 keV datasets. Interestingly, in the Alpine/120 keV data set, we observed a 2D class average for a partially dissociated LKB1 complex, something that we never have observed before in either data sets in this manuscript or previous internal work (Supplemental Fig. 3). Given that the samples were all prepared at the same time and in the same manner, we attribute this novel 2D class to the fact that extra contrast at 120 keV allows for better classification than previously possible.

To compare Gatan Alpine’s performance to the best imaging setup available to us, we also imaged the LKB1 complex with a TFS Titan Krios G2 at 300 keV with a BioQuantum energy filter and a K3 camera. Again, we started with a comparable initial number of particles for image processing and reached a similar final particle stack size (Supplemental Table 1). This yielded a reconstruction at 2.86 Å which was the highest resolution reconstruction out of all LKB1 complex datasets (Fig. 3D, Supplemental Fig. 3B). Interestingly however, the Alpine 120 keV dataset had the second highest resolution reconstruction of all 4 datasets, being bested only by the “gold-standard” microscope/detector configuration of a 3-condenser microscope with an energy filter and K3 detector.

As this is the first high resolution cryo-EM structure for the LKB1 complex, we compared it to the published crystal structure of the full complex (PDB: 2WTK) and the STRADα-MO25α subcomplex (PDB: 3GNI). Overall, the global conformation is similar to the crystal structure even though no nucleotide was added during sample preparation and we observed an apo LKB1 and an ADP-bound STRADα (Supplemental figure 4). Additionally, we were able to build some previously unresolved flexible loops and also observed some missing regions of the heterotrimeric complex in comparison to the crystal structure (Supplementary figure 5). Specifically, no density was observed for the STRADα Trp-Glu-Phe motif within the MO25α hydrophobic pocket although it was present in the construct and has been previously observed in the crystal structure. This motif is known to be required for MO25α recognition and binding in the absence of LKB1 (Boudeau et al. 2003). Another region not resolved in our structure as compared to the crystal structure was the c-terminus of STRADα (residues 401-414). This may be due to the fact that it packs against next molecule in the asymmetric unit in the crystal, packing not available in single particle cryo-EM (Supplementary figure 5B). Additionally, we resolved a more complete LKB1 C-terminal flanking tail (CTF) (residues 311-347) at the LKB1-STRADα interface resulting in novel interactions with the STRADα. (Supplemental figure 5). It remains to be investigated if these differences are of biological significance or are purely associated with differing methodologies utilized.

### 3.4 Using per-particle SSNR as a better metric for image quality

To compare the resulting reconstructions for both samples in a more quantitative manner, we compared the performance of each imaging system through per-particle spectral signal to noise (ppSSNR) plots. This analysis should be independent of the final particle number in reconstructions and reports on the inherent signal to noise ratio within each particle image. From this analysis, across different spatial frequencies, we observe that for the apoferritin datasets, the Alpine and K3 detectors perform similarly across spatial frequencies at the tested accelerating voltages until about 3.3 Å (Figure 4A, Supplemental figure 6). At higher spatial frequencies (up to resolution where FSC for each reconstruction crosses 0.143), we see that Alpine data set collected on TFS Glacios operating at 200 keV performs nearly identically if not better to the TFS Krios with Gatan K3 operating at 300 keV. We note again that the K3 datasets were collected on standard holey carbon grids, while the Alpine datasets were collected on UltrAuFoil grids, which have been shown to reduce beam-induced motion and enable higher resolutions than carbon (Russo and Passmore 2014). We observed somewhat different trends with the LKB1 complex ppSSNR curves. At lower frequencies (below 10 Å) all the imaging setups perform similarly, with the Alpine/120 keV slightly outperforming the K3/300 keV Krios and both of the 200 keV data sets. However, at higher frequencies (better than 5 Å) we observed substantial differences in ppSSNR, which followed the trends observed in the local resolution and Q-scores, wherein the Krios/K3/300 keV was the best, followed by Glacios/Alpine/120 keV, Glacios/K3/200 keV, and Glacios/Alpine/200 keV.

**Figure 4:**
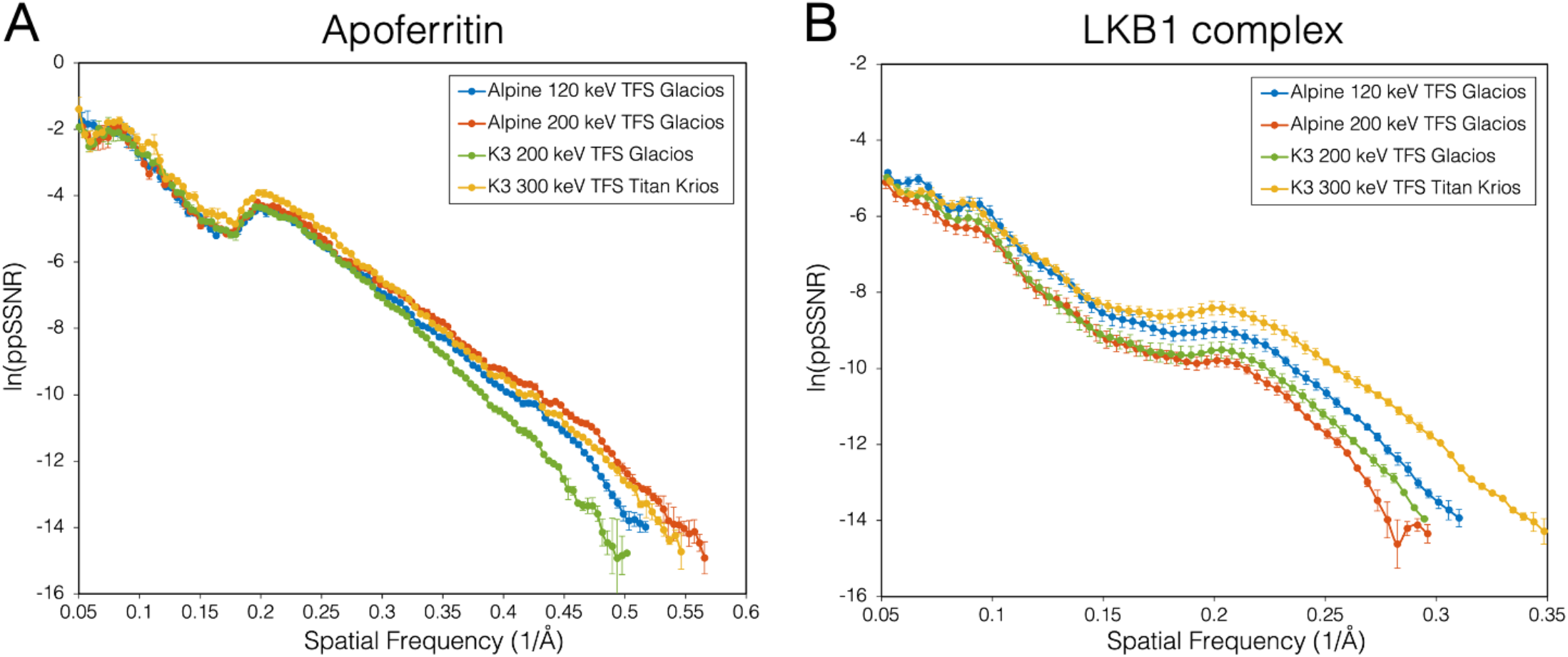
Comparison of data collections using per-particle SSNR. A plot of ppSSNR as a function of spatial frequency for (A) apoferritin and (B) LKB1 collected using the Alpine detector at 120 keV (blue), 200 keV (red), and the K3 detector at 200 keV (green) and 300 keV (yellow). Each curve is truncated to where the FSC from the smallest particle stack reaches 0.143.

### 3.5 Data collection at 100 keV on a Talos F200C with a side-entry cryo holder

Given the capacity of the Alpine detector to produce high-resolution structures on high-end microscopes that were equipped with an autoloader device, we next tested the utility of the Alpine detector in generating high-resolution data on a side-entry cryo-EM microscope, the TFS Talos F200C, at 200 keV and 100 keV. We began by imaging a standard sample apoferritin at 200 keV, an acceleration voltage that is typically used on this lens series to obtain high resolution structures when mounted with an autoloader (i.e. the Arctica and Glacios). Using a Gatan 626 side-entry cryo holder, an apoferritin grid was imaged after the sample had stabilized with a drift rate of less than 1 Å / sec. A small dataset of ~100 images with was acquired over the course of an hour, which was processed in RELION to yield a 2.7 Å resolution reconstruction from 21,174 particles (Supplemental figure 7, Supplementary table 4). Having confirmed that the Alpine detector could resolve a biological sample to better than 3 Å resolution at 200 keV with a side-entry holder, we next aligned the Talos F200C at 100 kV and loaded an apoferritin grid with the side-entry cryo holder for data acquisition. Using a similar number of particles (21,000), we were able to obtain a 3.1 Å resolution structure, and a little more than 2.5 times the amount of data (51,253 particles from 310 micrographs) was required to obtain a reconstruction at a comparable resolution of 2.6 Å resolution (Figure 5A, Supplemental figure 8A, Supplementary table 4).

We next tested the ability of this system to resolve a complex that was smaller than 200 kDa, so we targeted rabbit muscle aldolase, a well-characterized D2-symmetric tetramer that was previously determined to better than 3 Å resolution using 200 keV systems (Herzik, Wu, and Lander 2017). A little over 2,000 micrographs were acquired over the course of 1.5 days, refilling the side-entry holder every 1.5 hours, which were processed using standard image analysis routines. A final stack of 674,124 particles contributed to a ~3.1 Å resolution reconstruction, with regions resolved to better than 3 Å resolution according to the local resolution analysis (Figure 5B, Supplemental figure 8B, Supplementary table 4). Although the data collection is more labor-intensive (particularly overnight) than with an autoloader system, these findings demonstrate that high-resolution structures of smaller complexes can be determined with the Alpine camera using a side-entry holder at 100 keV.

**Figure 5:**
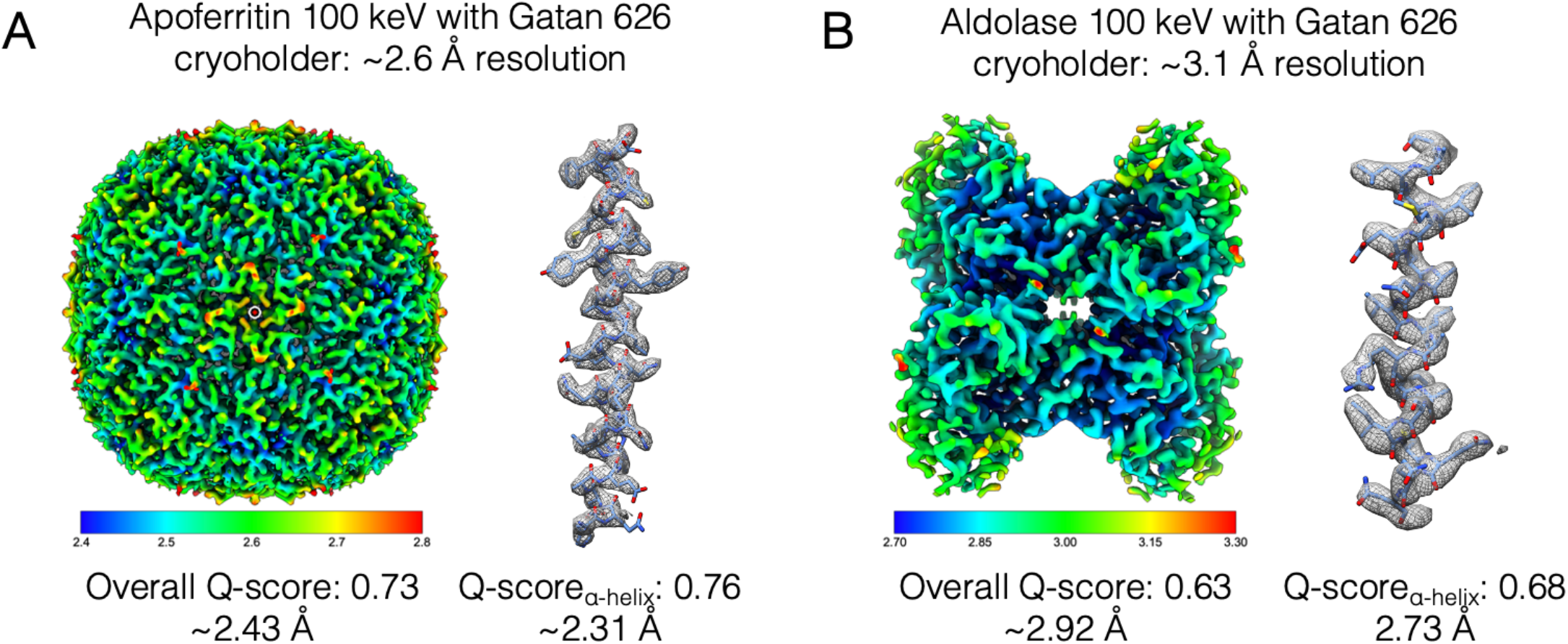
Reconstructions of apoferritin and aldolase using the Alpine detector on a Talos F200C microscope with a side-entry cryo-holder. 3D reconstructions of (A) apoferritin and (B) aldolase are colored by local resolution. To the right of each structure is a close-up view for a representative α-helix fit into each density map (apoferritin residues 13-42, aldolase residues 154-180). Overall Q-scores and Q-score derived effective resolution are shown below each structure and the representative helix.

Based on the success of our aldolase imaging, we next imaged much smaller complex – transythretin, which is a D2-symmetric tetramer amassing 55 kDa. This tetramer, which is prepared on grids containing a monolayer of graphene to overcome preferred orientation issues due to interactions with the air-water interface, was recently determined to better than 3 Å on a 200 keV system (Basanta et al. 2024), making this sample a viable candidate for imaging with the Alpine detector. A dataset of 7,001 movies of transthyretin on graphene were collected, but we were unable to determine a high-resolution structure of the tetramer. As discussed in the Methods, a workaround in the MotionCor2 program was used to process the 100 keV movies, thus it is possible that optimization of the dose-weighting curves applied frames could improve the quality of results. Notably, however, analysis of the first 300 micrographs during data acquisition yielded detailed 2D averages where secondary structure was clearly discernible (Supplemental figure 9). These 2D averages confirmed that the sample was sufficiently well-behaved and well-ordered for structural investigation, evenly distributed in the ice and readily amenable for particle picking, and that the particles adopted a range of different orientations on the grid. This type of information is important during the sample optimization process and could be gathered with only a few hours of data collection. Although a high-resolution 3D structure was not determined, these data highlight that imaging with an Alpine detector at 100 keV can inform on cryo-EM sample quality and viability for high-resolution structure determination on higher-end microscopes. Such a setup would have great utility at institutes that have side-entry microscopes with field emission guns, but whose projects are delayed due to a lack of screening access at national centers.

## 4. DISCUSSION

Our analyses show that the Gatan Alpine direct detector performs remarkably well in the context of multiple high-resolution cryo-EM imaging systems. We were able to obtain high resolution reconstructions of both apoferritin and a non-standard specimen, LKB1 heterotrimeric complex, illustrating the practical application of imaging at lower accelerating voltages. We were able to obtain a 1.76 Å reconstruction of apoferritin on UltrAuFoil grids from Alpine at 200 keV nearly reaching the quality of the best to date apoferritin reconstruction from the UCSF imaging center on carbon grids at 300 keV/K3/GIF and obtained with considerably more processing (1.65 Å). Excitedly, for LKB1 complex, Gatan Alpine at 120 keV performed better than both the Gatan K3 at 200 keV and itself at 200 keV, which we take as compelling evidence that Gatan Alpine is likely capturing the advantages of lower electron energy cryo-EM imaging, benefitting from the increased ratio of elastic to inelastic cross sections at 120 keV (Peet, Henderson, and Russo 2019).

It is important to note that our K3 datasets collected on the TFS Krios and TFS Glacios were collected in non-CDS mode, while all the Alpine data sets were collected in CDS mode. Although this may complicate direct comparison of the detectors, the UCSF facility and many other facilities operate their detectors in non-CDS mode for improved throughput foregoing the slight image quality improvement offered in CDS mode. Therefore, the data presented here provides a comparison between Alpine and the K3 in the mode it is most likely to be operated in. Additionally, previous work demonstrated very small differences between CDS and non-CDS modes for particles comparable in size to the LKB1 complex (aldolase, (Sun et al. 2021)). Therefore, although we cannot rule out if the K3 operated in CDS mode at 200 keV would outperform Alpine, it is unlikely.

For the LKB1 complex sample, Gatan K3 at 300 keV on a TFS Titan Krios with BioQuantum energy filter, which we have used to represent the best state-of-the-field cryo-EM imaging setup still outperformed Glacios/Alpine data set at 120 keV. This may be due the recently reported observation that the key limitation to low energy imaging is the chromatic aberration coefficient, or *C*_*c*_ (Naydenova et al. 2019; McMullan et al. 2023). It is worth noting that our experiments used a specially-built TFS Glacios with a uniquely narrow pole gap of 7 mm, called the SP-Twin lens yielding the *C*_*c*_ of 1.7 mm and a *C*_*s*_ of 1.6 mm (TFS, direct communication), potentially ameliorating this effect. Several hardware improvements have been proposed to reduce the *C*_*c*_ limitation, such as better designed objective lenses and dedicated *C*_*c*_ correctors. With those imaging setups operating at 120 keV in conjunction with the Gatan Alpine detector, it is possible that the better signal to noise yielded at lower voltages will be captured, providing an optimal imaging set up surpassing current state of the art systems.

Still, the Gatan Alpine alone exhibits huge improvements to the feasibility of low energy ~100 keV cryo-EM imaging, outperforming the Gatan K3 at 200 keV, where we would expect it to perform even worse at ~100 keV. Even despite not yet achieving greater resolutions than state-of-the-art imaging systems, Gatan Alpine still represents a great alternative for microscopes operating at voltages of 100-200 keV which are often more accessible than the state of the art 300 keV systems. Furthermore, we expect lower energy cryo-EM imaging, with improved contrast, to be particularly advantageous for smaller biological samples.

The exponentially rising demand for cryo-EM across the biology community, and the need to drive down costs of both data collection and screening, requires a high speed, high DQE, large-format direct electron detector optimized for lower accelerating voltages. At the same time, lower energy cryo-EM imaging will also provide the inherent benefits of improved radiation damage and image contrast. Both the democratization of cryo-EM, as well as the drive for optimal cryo-EM imaging, can be simultaneously answered with a new direct electron detector. The camera presented here fills these needs, and will pave a path towards both a more widespread use of cryo-EM imaging, and the imaging of smaller samples by cryo-EM.

## ABBREVIATIONS (footnote)

Cryo-EM: Cryo-Electron Microscopy
DQE: Detective Quantum Efficiency
CDS: Correlated Double Sampling
FSC: Fourier Shell Correlation
kV: kilovolt
keV: kiloelectron volt
LKB1: Liver kinase B1
STRADα: STE20-related kinase adapter protein alpha
MO25α: Mouse protein 25 alpha
TTR: Transthyretin

## OTHER INFO

### 1. CRediT AUTHORSHIP CONTRIBUTION STATEMENT

**Lieza Chan**: Conceptualization, Data curation, Formal analysis, Investigation, Visualization, Writing - original draft, Writing – review & editing. **Brandon Courteau**: Conceptualization, Data curation, Formal analysis, Investigation, Visualization, Writing - original draft, Writing – review & editing. **Allison Maker:** Conceptualization, Data curation, Formal analysis, Investigation, Visualization, Writing - original draft, Writing – review & editing. **Mengyu Wu:** Data curation, formal analysis, investigation, methodology. **Benjamin Basanta:** Formal analysis, investigation. **Hevatib Mehmood**: Resources. **David Bulkley**: Conceptualization, Formal Analysis, Investigation, Methodology, Resources. **David Joyce:** Data curation, Writing – review & editing. **Brian C. Lee**: Data curation, Writing – review & editing. **Stephen Mick**: Project administration, Resources, Supervision, Writing – review & editing. **Sahil Gulati**: Project administration, Methodology, Resources, Supervision, Writing – original, Writing – review & editing. **Gabriel C. Lander**: Conceptualization, Data curation, Formal analysis, Funding acquisition, Investigation, Methodology, Project administration, Resources, Software, Supervision, Validation, Visualization, Roles/Writing - original draft, and Writing - review & editing. **Kliment A Verba:** Conceptualization, Funding acquisition, Investigation, Methodology, Project administration, Resources, Software, Supervision, Validation, Visualization, Roles/Writing - original draft, and Writing - review & editing.

## 3. ACKNOWLEDGEMENTS

We thank Glenn Gilbert (UCSF) for help with microscope operations, Matt Harrington (UCSF) for microscope computational support, Shawn Zheng (UCSF) for help with finding optimal parameters for MotionCorr2, Gary Chan (UCSF) and John Gordan (UCSF) for LKB1 complex plasmid. We thank David Agard, Yifan Cheng and broader UCSF CryoEM supergroup for helpful comments regarding the project. We thank Bill Anderson (Scripps Research) for 100 keV microscope alignment support, Charles Bowman (Scripps Research) for help integrating the Alpine detector for data collection, and Jean-Christophe Ducom (Scripps Research High Performance Computing) for computational support. We thank Christopher R. Booth and Paul Mooney (Gatan Inc., Pleasanton, CA) for helpful comments.

