## Supplemental Material for "High-resolution single-particle imaging at 100-200 keV with the Gatan Alpine direct electron detector"

### SUPPLEMENTAL INFORMATION

**Table S1.** Data collection parameters for each collection

| | Magnification (nominal/ calibrated) | Total exp ($e^-$/Å$^2$) | Exposure | | Exp rate ($e^-$/pix/s) | Mode | Pixel size (Å/physpix) | MotionCorr parms (v1.4.0) |
|---|---|---|---|---|---|---|---|---|
| Alpine 120 keV TFS Glacios | 43,000 / 58,823 | 58.9 | 0.595 $e^-$/A$^2$/frame | 0.05s frame | 8.6 | CDS | 0.85 | 3x4 patches Group 3 frames |
| Alpine 200 keV TFS Glacios | 43,000 / 54,229 | 45.5 | 0.46 $e^-$/A$^2$/frame | 0.05s frame | 7.8 | CDS | 0.922 | 3x4 patches Group 3 frames |
| K3 200 keV TFS Glacios | 54,000 / 68,493 | 55.1 | 0.60 $e^-$/Å$^2$/frame | 0.02s frame | 16 | nonCDS | 0.73 | 7x5 patches Group 1 frame |
| K3 300 keV Titan Krios1 (LKB1) | 105,000 / 59,880 | 45.8 | 0.57 $e^-$/Å$^2$/frame | 0.025s frame | 16 | nonCDS | 0.835 | 7x5 patches Group 1 frame |
| K3 300 keV Titan Krios1 (apoferritin) | 105,000 / 59,880 | 47.7 | 0.596 $e^-$/Å$^2$/frame | 0.025s frame | 16 | nonCDS | 0.8189 | 7x5 patches Group 1 frame |
| Alpine 200 keV Talos (apoferritin) | 45,000 / 57,339 | 51.4 | 1.02 $e^-$/Å$^2$/frame | 0.1s frame | 7.8 | CDS | 0.872 | 3x4 patches Group 1 frames |
| Alpine 100 keV Talos (apoferritin) | 45,000 / 58,685 | 143.3 | 0.72 $e^-$/Å$^2$/frame | 0.1s frame | 5.2 | CDS | 0.852 | 3x4 patches Group 1 frames |
| Alpine 100 keV Talos (aldolase) | 45,000 / 58,685 | 71.6 | 0.72 $e^-$/Å$^2$/frame | 0.1s frame | 5.2 | CDS | 0.852 | 3x4 patches Group 1 frames |
| Alpine 100 keV Talos (transthyretin) | 45,000 / 58,685 | 50.1 | 0.72 $e^-$/Å$^2$/frame | 0.1s frame | 5.2 | CDS | 0.852 | 3x4 patches Group 1 frames |

**Table S2.** Comparison of image processing and reconstructions for apoferritin TFS Glacios and Titan Krios collections

| | Initial micrographs (no.) | Curated micrographs (no.) | Initial particles (no.) | Final particles (no.) | NU-Refinement | |
|---|---|---|---|---|---|---|
| | | | | | Res (Å) | B-factor |
| Alpine 120 keV TFS Glacios | 3,789 | 3,312 | 1,635,971 | 200,000 | 1.92 | 65.2 |
| Alpine 200 keV TFS Glacios | 5,046 | 3,619 | 1,834,404 | 204,387 | 1.76 | 57.7 |
| K3 200 keV TFS Glacios | 5,097 | 3,903 | 2,675,691 | 200,000 | 1.98 | 79.7 |
| K3 300 keV Titan Krios1 | 622 | 617 | 790,751 | 200,000 | 1.82 | 65.4 |

**Table S3.** Comparison of image processing and reconstructions for LKB1 complex TFS Glacios and Titan Krios collections

| | Initial micrographs (no.) | Curated micrographs (no.) | Initial particles (no.) | Final particles (no.) | NU-Refinement | |
|---|---|---|---|---|---|---|
| | | | | | Res (Å) | B-factor |
| Alpine 120 keV TFS Glacios | 3,635 | 2,868 | 897,749 | 120,684 | 3.19 | 152.3 |
| Alpine 200 keV TFS Glacios | 3,635 | 3,179 | 1,229,097 | 111,117 | 3.37 | 163.2 |
| K3 200 keV TFS Glacios | 1,504 | 1,337 | 1,000,313 | 144,801 | 3.37 | 177.1 |
| K3 300 keV Titan Krios1 | 1,293 | 1,223 | 979,927 | 124,780 | 2.86 | 121.1 |

**Table S4.** Comparison of image processing and reconstructions for Talos F200C collections

| | Scope/ detector setup | Initial micrograph s (no.) | Curated micrograph s (no.) | Initial particles (no.) | Final particles (no.) | NU-Refinement | |
|---|---|---|---|---|---|---|---|
| | | | | | | Res (Å) | B-factor |
| apoferritin | Alpine 100 keV Talos | 310 | 218 | 82,977 | 59,000 | 2.60 | 103.4 |
| | Alpine 200 keV Talos | 100 | 84 | 29,404 | 16,000 | 2.70 | 90.9 |
| aldolase | Alpine 100 keV Talos | 2,476 | 2,027 | 1,914,721 | 674,124 | 3.09 | 189.2 |

**Table S5.** Statistics of data collection, cryo-EM refinement, and the resulting atomic model for LKB1 complex collected at 8eps and 105kx in CDS mode (Relevant to Figure 3) on TFS Krios.

| LKB1 complex (K3/Krios) | |
|---|---|
| Data collection and processing | |
| Magnification | 105,000 |
| Voltage (kV) | 300 |
| Electron exposure ($e^-/Å^2$) | 69 |
| Defocus range (µm) | -1.0 to -2.0 |
| Physical pixel size (Å) | 0.835 |
| Map resolution (Å) | 2.86 (FSC 0.143) |
| Model Refinement | |
| Initial model used (PDB code) | 2WTK |
| Model resolution (Å)<br>(FSC threshold 0.143/0.5)<br>CC volume/mask | 2.8/3.1 (0.143/0.5)<br>0.83/0.87 |
| Model composition<br>Atoms non-H<br>Protein residues<br>Ligands | <br>7513<br>930<br>1 |
| B factors ($Å^2$) min/max/mean<br>Protein<br>Ligand | <br>13.02/99.40/39.00<br>30.43 |
| R.m.s. deviations<br>Bond lengths (Å) (#>4σ)<br>Bond angles (°) (#>4σ) | <br>0.012<br>1.483 |
| Validation<br>MolProbity score<br>Clashscore<br>Poor rotamers (%)<br>CaBLAM outliers (%)<br>Q-score | <br>0.70<br>0.60<br>0<br>0.23<br>0.66 |
| Ramachandran plot<br>Favored (%)<br>Allowed (%)<br>Disallowed (%) | <br>98.70<br>1.30<br>0.00 |
| Data Availability | |
| EMDB<br>PDB<br>EMPIAR | EMD-43506<br>PDB: 8VSU |

**Supplemental Figure 1:** Gold-standard and directional FSC curves for the apoferritin reconstructions from data collections collected with the Alpine detector with the TFS Glacios at 120 keV and 200 keV, and the K3 detector on the TFS Glacios at 200 keV and the TFS Titan Krios at 300 keV.

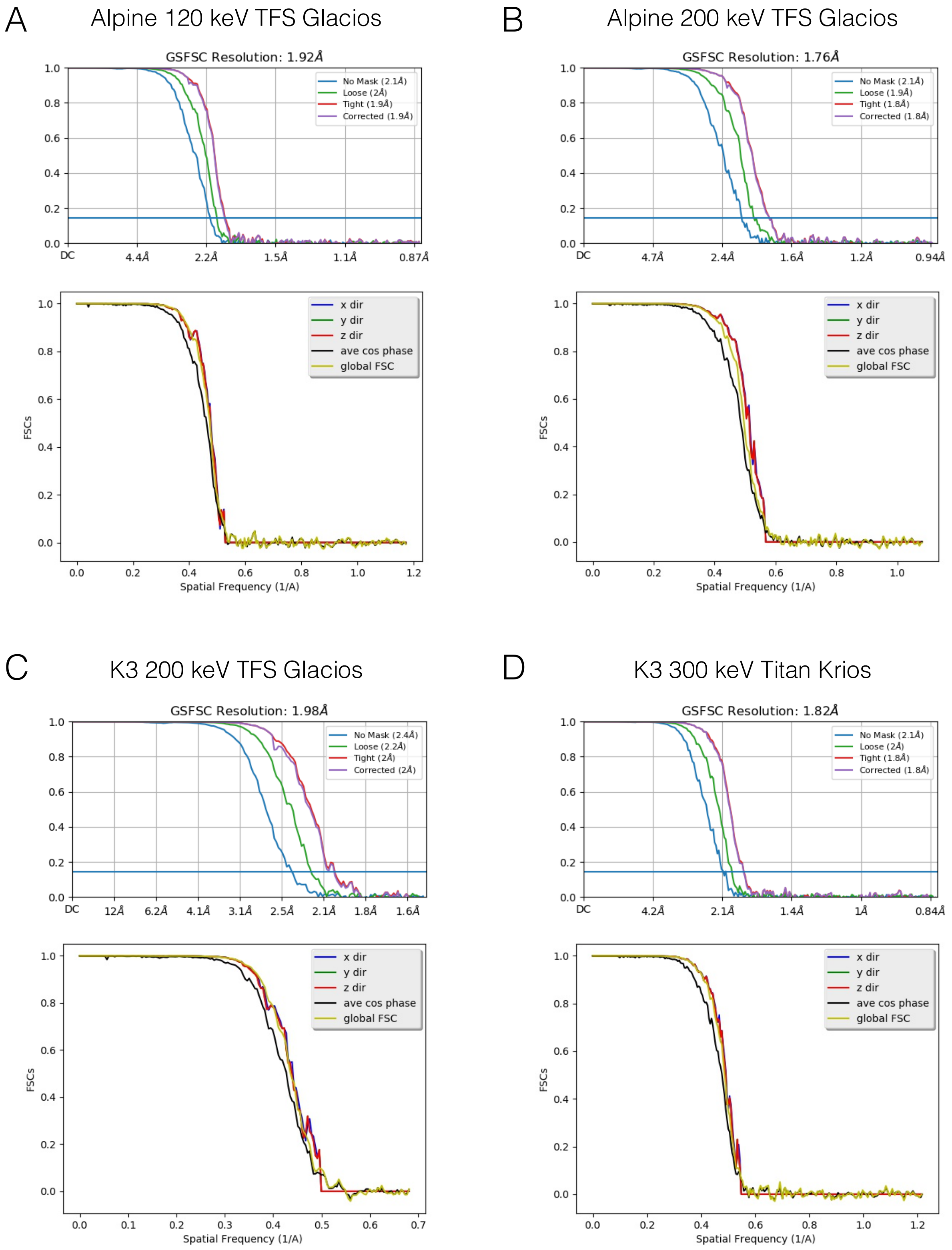

**Supplemental Figure 2: The LKB1 complex imaging and reconstruction data for Alpine detector at 120 keV and 200 keV. (**A) Schematic of LKB1 heterocomplex and subcomplex. LKB1 shown in pink, MO25α shown in blue, and STRADα shown in green. Representative micrograph, 2D class averages, gold-standard FSC curves, and directional FSC curves for all data collections collected with the Alpine detector with the TFS Glacios at (B) 120 keV and (C) 200 keV. Novel 2D class average for subcomplex is demarcated by purple border.

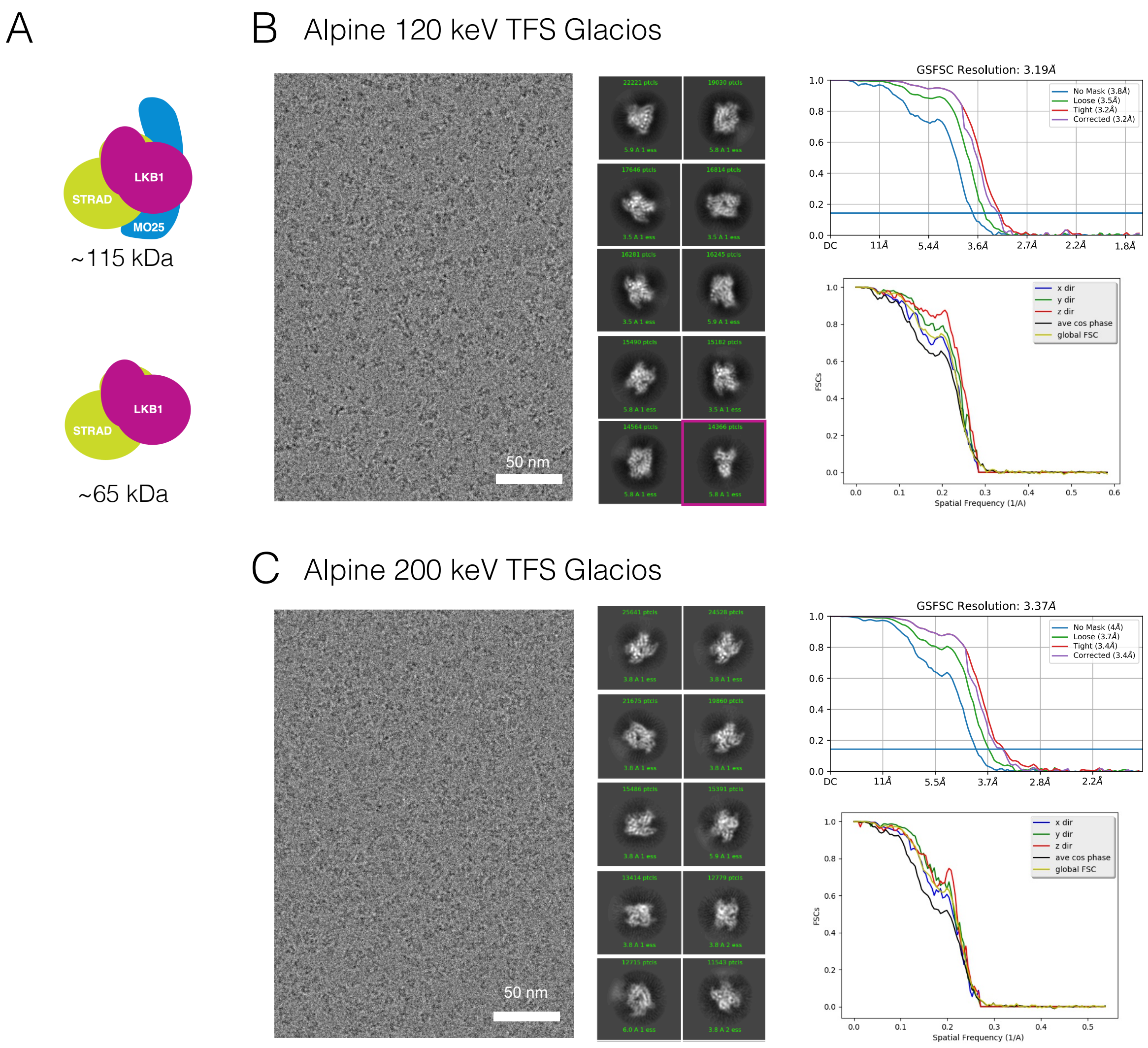

**Supplemental Figure 3: The LKB1 complex imaging and reconstruction for K3 detector collections.** Representative micrograph, 2D class averages, gold-standard FSC curves, and directional FSC curves for data collections with the K3 detector on the (A) TFS Glacios at 200 keV and the (B) TFS Titan Krios at 300 keV.

A K3 200 keV TFS Glacios

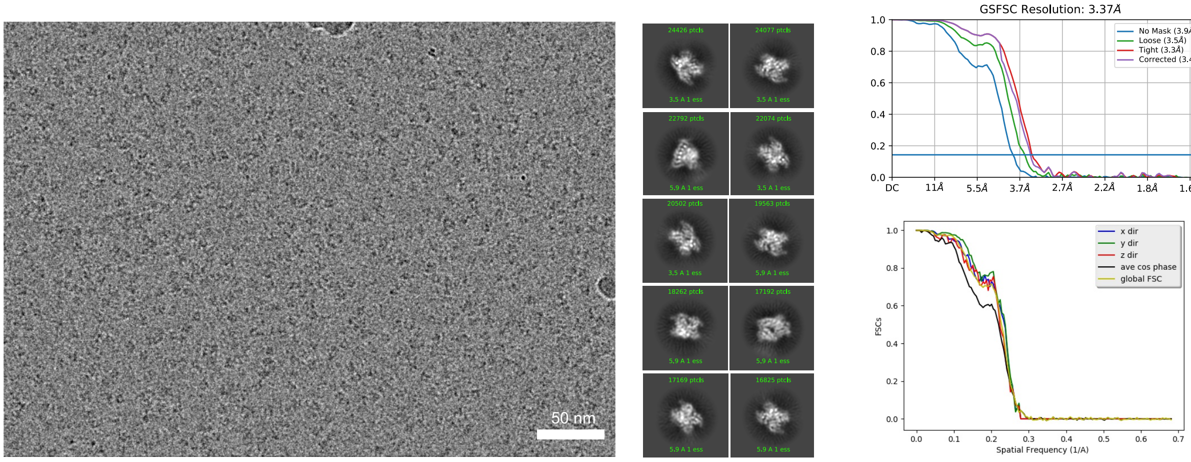


B K3 300 keV Titan Krios

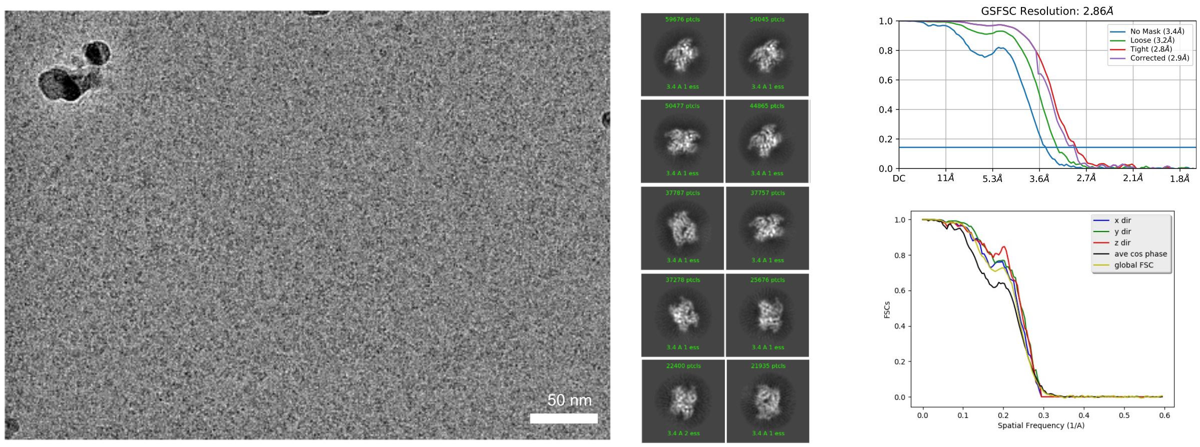

**Supplemental Figure 4: LKB1-STRADα-MO25α heterocomplex is in a different nucleotide state in cryo-EM reconstruction as compared to the previously published crystal structure.** A) Cryo-EM map and the resulting model of the LKB1 heterocomplex with LKB1 shown in pink, MO25α shown in blue, and STRADα shown in green. B) Overlay of the cryo-EM built model in ribbon (colored) and the crystal structure of the LKB1 complex (PDB: 2WTK) in grey. C) Zoomed-in view showing cryo-EM density of the nucleotide pocket of the ADP-bound STRADα pseudokinase. D) Zoomed-in view showing cryo-EM density of the nucleotide pocket of the apo LKB1 kinase. AMPPNP from the crystal structure is shown to demonstrate lack of its cryo-EM density.

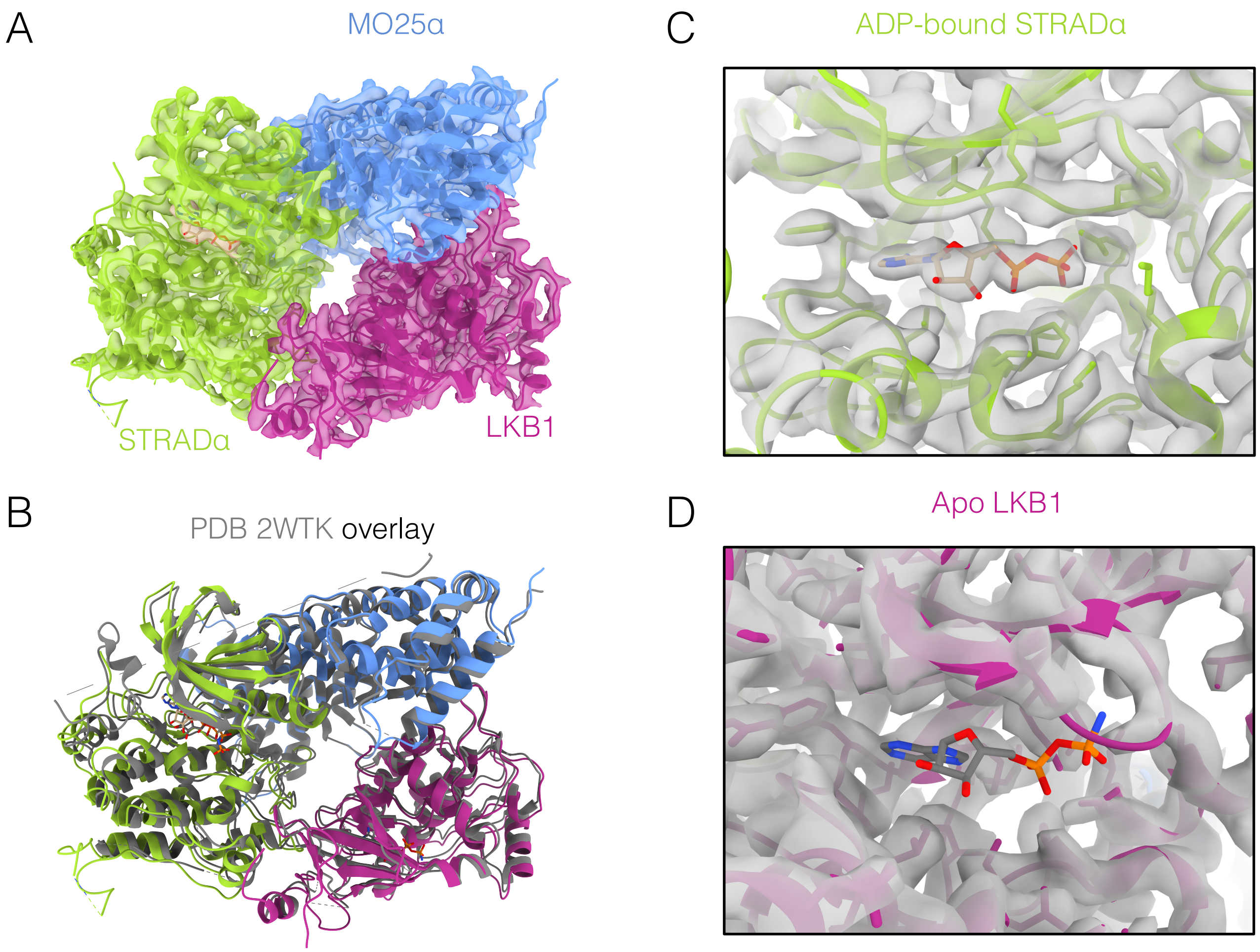

**Supplemental Figure 5: Notable differences between cryo-EM model and the crystal structure of LKB1-STRADα-MO25α heterocomplex.** A) Cryo-EM density and resulting model for MO25α (blue ribbon) with overlay of the STRADα WEF motif from the crystal structure in green showing no density for that motif. Two rotamer conformations of Mo25α ARG256 with EM density in light blue. B) Cryo-EM map (transparent gray) and model colored by protein (as in sup fig 4A). C terminal region of STRADα for which cryo-EM density is missing is outlined in red that and is at the crystal packing interface with the neighboring molecule shown in gray surface. C) Now resolved LKB1 C-terminal Flanking Tail (pink) overlayed with the crystal structure (gray). Dashed lines represent missing residues. Novel interactions at binding interfaces between LKB1 C-terminal tail and STRADα at (D) αG helix and (E) αEF-αF loop are displayed. D) Additional residues observed in the cryo-EM structure are shown in light pink. Interactions shown by dotted lines. E) EM density and fitted model of the C-terminal flanking tail interaction of TRP332 with GLN254 of the STRADα αEF-αF loop.

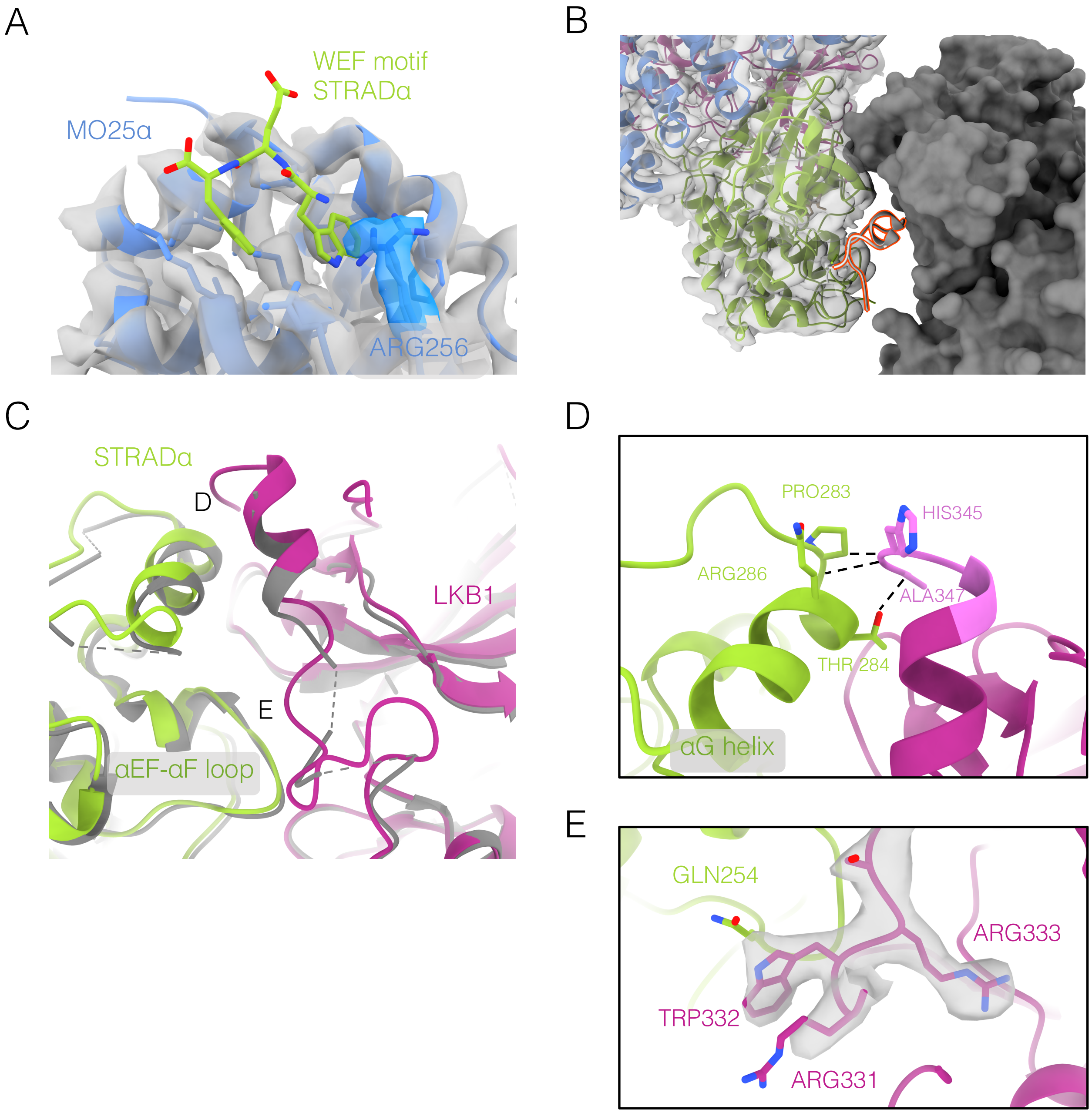

**Supplemental Figure 6: Comparison of image quality using per-particle SSNR in terms of fraction of Nyquist.** A plot of ppSSNR as a function of fraction of physical Nyquist for (A) apoferritin and (B) LKB1 collected using the Alpine detector at 120 keV (blue), 200 keV (red), and the K3 detector at 200 keV (green) and 300 keV (yellow). See Table S1 for explicit pixel sizes associated with these collections.

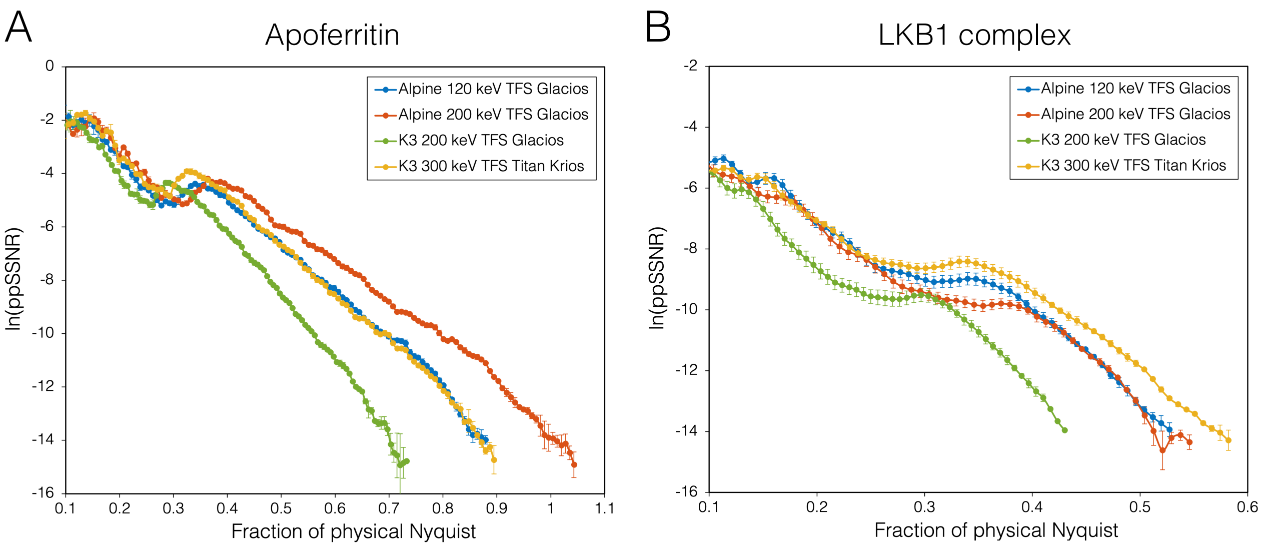

**Supplemental Figure 7: 200 keV apoferritin reconstruction using the Alpine detector on a Talos F200C microscope with a side-entry cryo-holder.** 3D reconstruction of apoferritin colored by local resolution (left), and the gold-standard global FSC curve and the directional FSC curves (right).

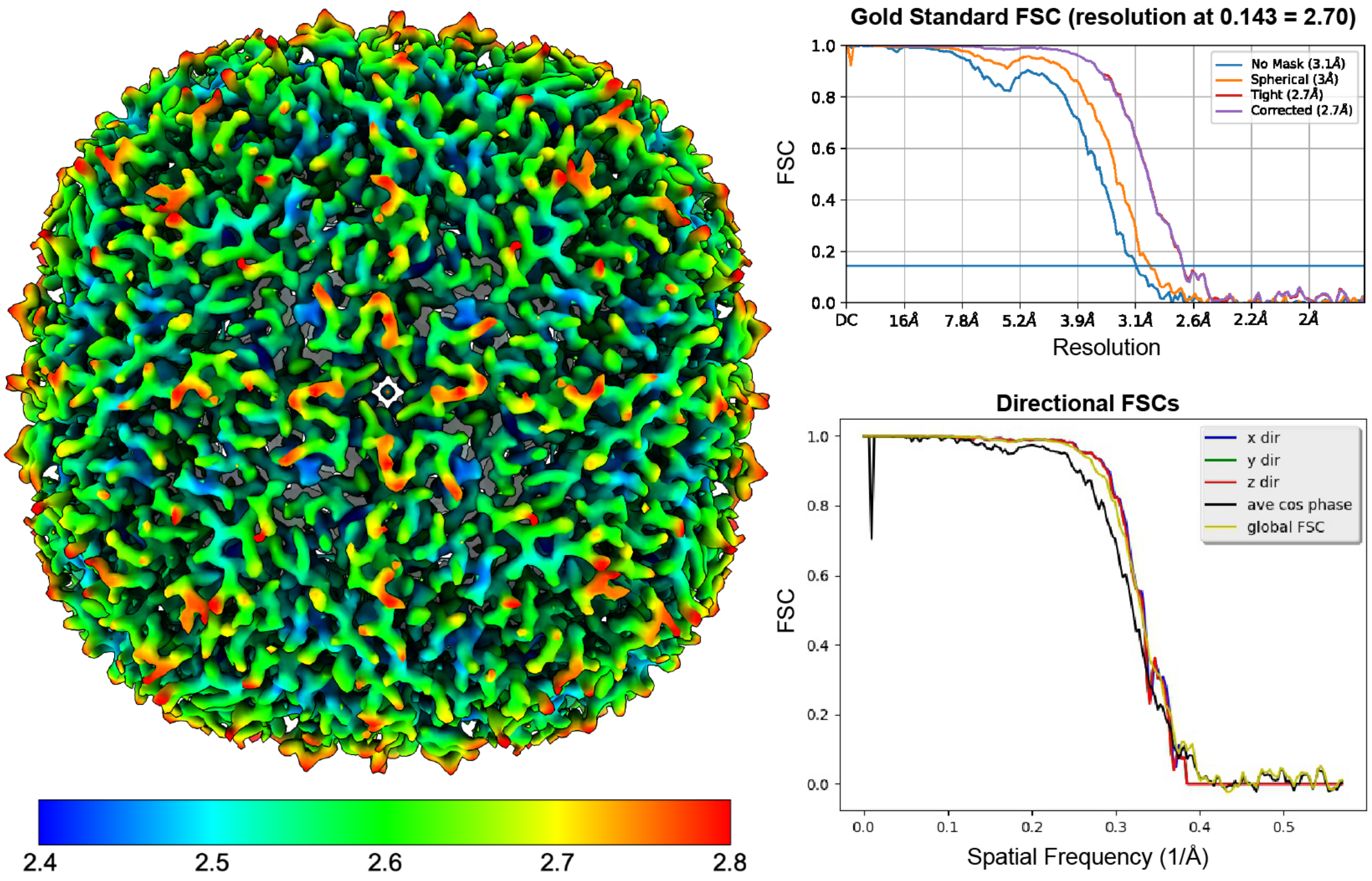

**Supplemental Figure 8: 100 keV imaging and reconstruction with the Alpine detector on a Talos F200C with a side-entry cryo-holder.** Representative micrograph, gold-standard FSC curves, and viewing direction distribution for (A) apoferritin and (B) aldolase samples.

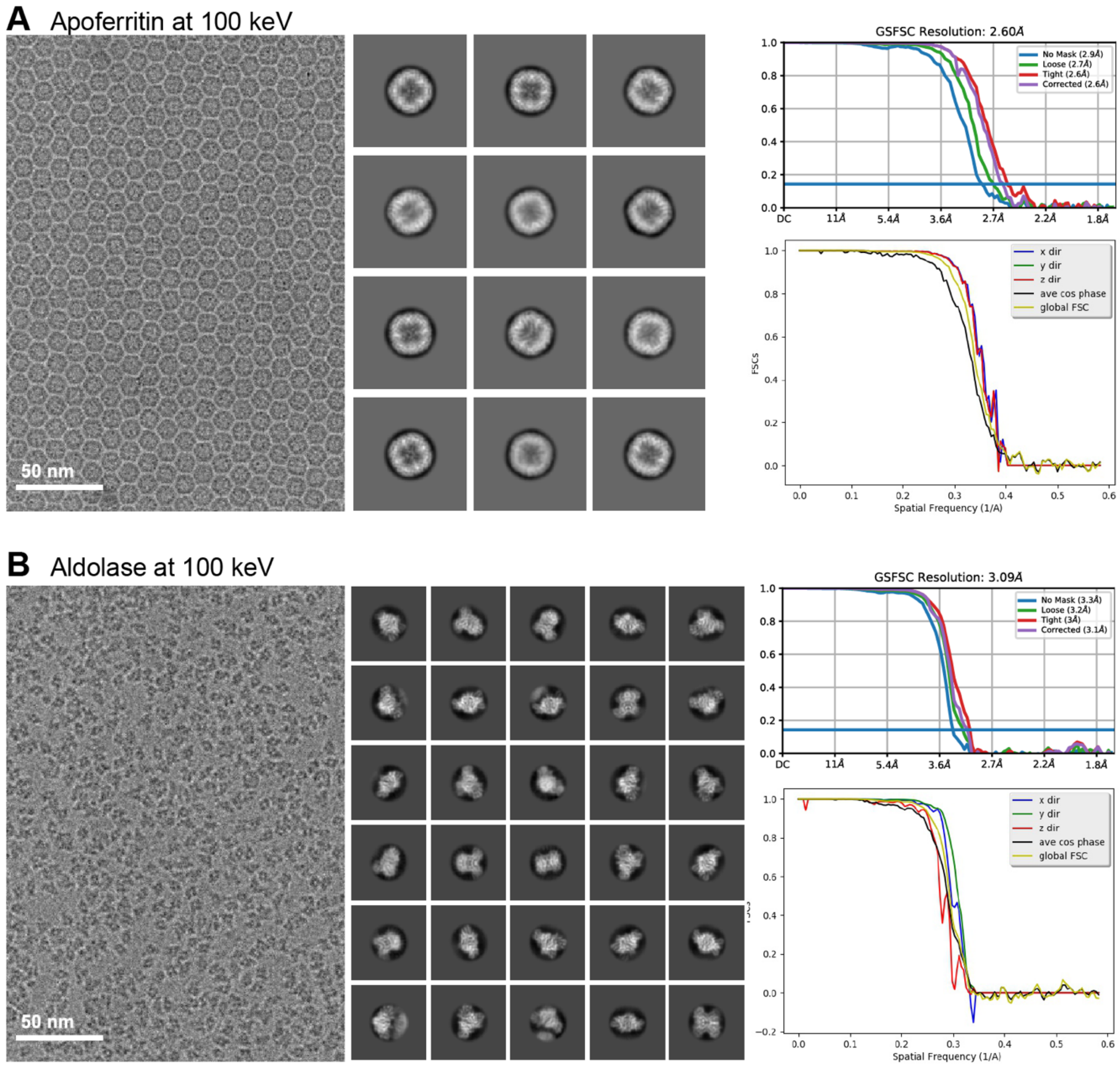

**Supplemental Figure 9**: 100 keV imaging of transthyretin (55 kDa) on graphene. Representative micrograph (left) and 2D class averages (right) of transthyretin sample imaged with the Alpine detector on a Talos F200C at 100 keV with a side-entry cryo-holder. Secondary structure is clearly visible in the 2D averages.

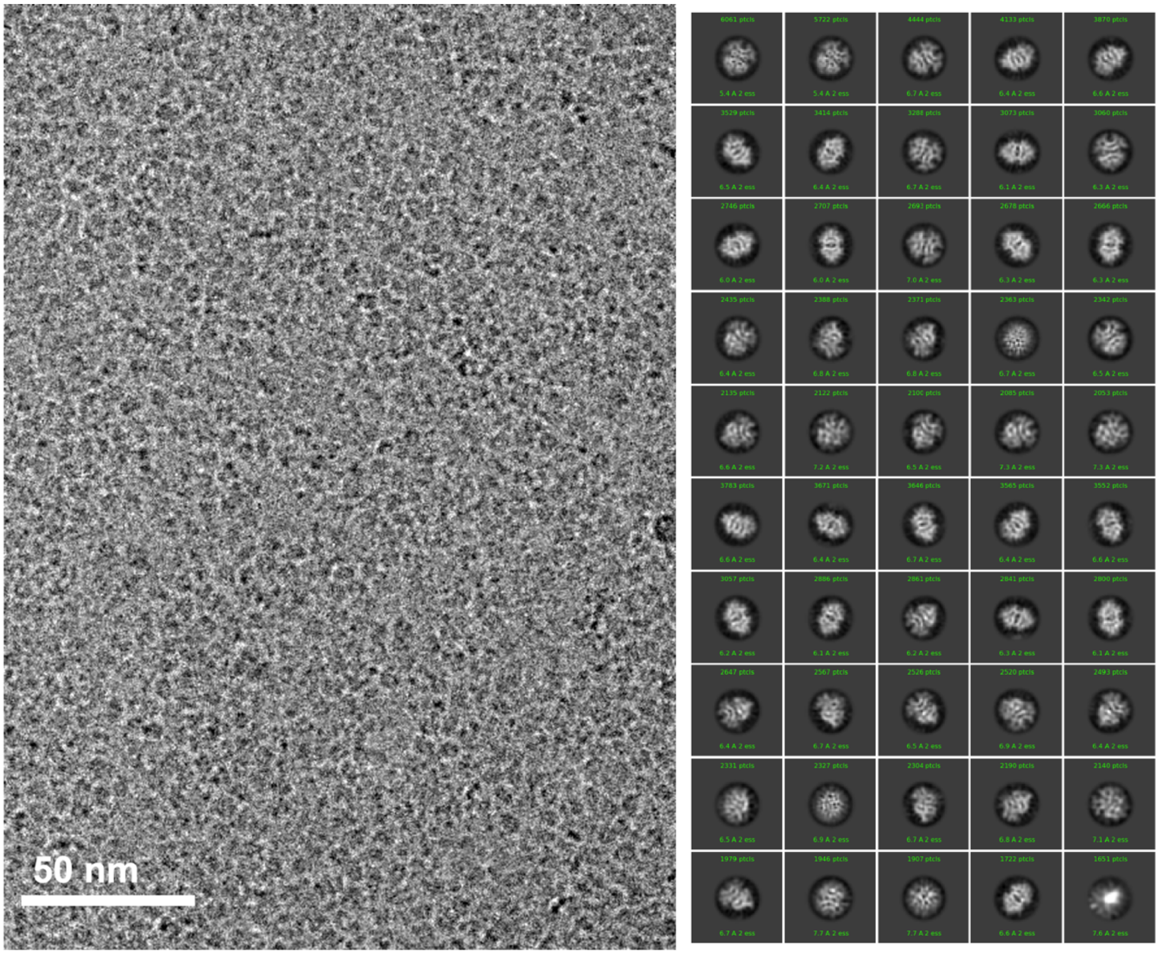
